# Molecular details of the CPSF73-CPSF100 C-terminal heterodimer and interaction with Symplekin

**DOI:** 10.1101/2023.04.19.537554

**Authors:** Stéphane Thore, Finaritra Raoelijaona, Vincent Talenton, Sébastien Fribourg, Cameron D. Mackereth

## Abstract

Eukaryotic pre-mRNA is processed by a large multiprotein complex to accurately cleave the 3’ end, and to catalyze the addition of the poly(A) tail. Within this cleavage and polyadenylation specificity factor (CPSF) machinery, the CPSF73 endonuclease subunit directly contacts both CPSF100 and the scaffold protein Symplekin to form a subcomplex known as the core cleavage complex (CCC) or mammalian cleavage factor (mCF). Here we have taken advantage of a stable CPSF73-CPSF100 minimal heterodimer from E. cuniculi to determine the solution structure formed by the first and second C-terminal domain (CTD1 and CTD2) of both proteins. We find a large number of contacts between both proteins in the complex, and notably in the region between CTD1 and CTD2. A similarity is also observed between CTD2 and the TATA-box binding protein (TBP) domains. Separately, we have determined the structure of the terminal CTD3 domain of CPSF73, which also belongs to the TBP domain family and is connected by a flexible linker to the rest of CPSF73. Biochemical assays demonstrate a key role for the CTD3 of CPSF73 in binding Symplekin, and structural models of the trimeric complex from other species allow for comparative analysis and support an overall conserved architecture.

## INTRODUCTION

Pre-messenger RNA, with the notable exception of histone-coding pre-mRNA, requires processing at the 3’-end via sequential steps of cleavage and polyadenylation. The resulting polyadenosine tail-containing mRNA is protected from the nuclear degradation machinery and ready for cytoplasmic translation. The process of cleavage and polyadenylation is also subject to regulation, and for example the choice of polyadenylation signal (PAS) on the pre-mRNA among alternate sites is a regulatory aspect of cell-type function and development. Furthermore, perturbation of this process is known to severely impact cell physiology, with links to a broad range of cancers and disease (1).

The 3’-end processing of pre-mRNA in metazoa is performed by the large multi-protein Cleavage and Polyadenylation Specificity Factor (CPSF). Metazoan CPSF is composed of a seven subunit core, which can be divided into stable sub-complexes with specific functions. The sub-complex consisting of CPSF160, CPSF30, WDR33, and hFip1, is known in mammals as the mammalian polyadenylation specificity factor (mPSF) that binds the PAS motif and recruits poly(A) polymerase (2). The second sub-complex is required for the cleavage step, and is composed of CPSF73, CPSF100 and Symplekin. These three proteins are co-purified under stringent conditions, and their presence in both the canonical CPSF complex, as well as the complex required to process histone pre-mRNA, identify them as the essential core cleavage factors (3)(4). Based on later analyses, the sub-complex is also named the Core Cleavage Complex (CCC) common to all metazoan 3’-end processing complexes (5), or the mammalian Cleavage Factor (mCF) based on the human proteins (6).

The CPSF73 subunit is a member of the metallo β-lactamase (MBL) family of proteins (7), and is the catalytically active component in CCC/mCF (8)(9)(10)(11). Specifically, the N-terminus of CPSF73 has an MBL fold that contains the active site residues (12), and also a β-CASP insert which may assist in RNA binding. CPSF73 was shown to directly contact the cleavage site downstream of the AAUAAA motif on the pre-mRNA, and its activity is zinc-dependent (10). Additional point mutants demonstrated that CPSF73 is the endonuclease within the histone 3’ processing complex (12). The second subunit of CCC/mCF, CPSF100, shares a similar domain architecture with CPSF73 but does not exhibit endonuclease activity (11)(12). Nevertheless, the presence of CPSF100 is required to enable functional activity of CPSF73 and form a tether to the rest of the CPSF (10)(11)(12). The third protein, Symplekin, is a long HEAT repeat-containing protein acting as a hub for the interaction with multiple factors, and was identified as part of the CPSF from *Xenopus* studies (13), the mouse histone 3’ processing complex (14), and by comparison to the Pta1 subunit from the yeast Cleavage and Polyadenylation Factor (CPF) (15)(16).

The atomic details for several regions in CPSF73, CPSF100, and Symplekin, have been described based on isolated domains or within larger assemblies. The crystal structure of the N-terminal MBL region of human CPSF73 (hsCPSF73-MBL) revealed two zinc-binding sites, and allowed for structure-guided mutagenesis to confirm active site residues with comparison to the structure of yeast CPSF100 (Cft2) (11). Crystal structures of *Cryptosporidium* CPSF73-MBL were obtained in the apo form and bound to an anti-parasite compound (17), and hsCPSF73-MBL was crystallized in the presence of the anti-cancer drug JTE-607 (18). Structure of the N-terminal region of Symplekin was first determined from *Drosophila* (19), followed by the human N-terminal region in complex with Ssu72 and a RNAPII-CTD phosphopeptide (20)(21). Cryo-EM structures of human CPSF (6) and the histone pre-mRNA 3’-end processing complex (22) provided valuable high-resolution insight into the architecture of the N-terminal regions of CPSF73, CPSF100 and Symplekin within the larger processing machinery. In contrast, there was significantly lowered resolution observed for the C-terminal regions of the CCC/mCF proteins, which necessitated homology docking from the IntS11-IntS9 crystal structure (23), as well as poly-Ala helices to fit the Symplekin C-terminal density. Despite the limited resolution, it is clear that the C-terminal regions of CPSF73, CPSF100 and Symplekin come together to form the main contacts within the CCC/mCF. The histone pre-mRNA 3’-end processing complex structure further defined three C-terminal domains for CPSF73 (CTD1, CTD2, CTD3) and two C-terminal domains for CPSF100 (CTD1, CTD2) (22). There was evidence that the CTD1 from both CPSF73 and CPSF100 come together to form a single β-barrel, and that the CTD2 from each protein resembles the C-terminal heterodimer of IntS11-IntS9 (23). Additional density further suggested that CTD3 from CPSF73 may contact Symplekin. However, specific atomic information for these regions was not possible to determine.

The direct interaction between the C-termini of CPSF73, CPSF100, and Symplekin, have also been studied using biochemistry and molecular biology. Using *Drosophila* proteins, it was noted that the minimal regions required to form the CCC/mCF complex involved the C-termini of dmCPSF73 (residues 552-684) and dmCPSF100 (residues 648-756), and the dmSympk region from residue 272-1080 (5). In *Arabidopsis*, it was shown that the last 157 residues of atCPSF100 interact strongly with the last 254 residues of atCPSF73-I, or the last 180 residues of the truncated paralogue atCPSF73-II (24). Insight into the CCC/mCF C-terminal complex has also been provided from Integrator, in which the Integrator subunits IntS11, IntS9, and IntS4, are closely related to CPSF73, CPSF100, and Symplekin, respectively. The IntS11-IntS9-IntS4 trimer represents a minimal Integrator cleavage module, similar to that of CCC/mCF, and is stable during purification (25)(26). It had been noted previously that the C-termini of IntS11 and IntS9 strongly interact (27), and the crystal structure of a IntS11-IntS9 heterodimer of the CTD2 from each protein (23) was already helpful for analysis of the histone pre-mRNA 3’-end processing complex (22). Several recent cryo-EM structures containing Integrator have provided improved resolution for the CTD1 and CTD2 domains of IntS11 and IntS9 (26)(28)(29). Insight into the metazoan CPSF is also provided by structural comparison to the yeast Cleavage and Polyadenylation Factor (CPF) nuclease module which contains Ysh1 and Cft2 (orthologues to CPSF73 and CPSF100), but at present the C-termini of these two proteins have not been observable by structural methods (30)(31).

To obtain this missing atomic information on the assembly of the CCC/mCF we have taken advantage of the CPSF73 and CPSF100 proteins from *E. cuniculi* to determine the solution structure of all C-terminal domains. This includes an intimate complex formed by CTD1 and CTD2 of both proteins, as well as the independent CTD3 domain of CPSF73 that we find by biochemical analysis to be essential for binding Symplekin.

## MATERIALS AND METHODS

### Cloning

Plasmids expressing hsCPSF73, hsCPSF100, ecCPSF73, ecCPSF100 and ecSymplekin were built by inserting the PCR-amplified sequences into modified pET vectors allowing for coexpression with or without an N-terminal hexa-histidine Tag. The plasmids used for co-expression of ecCPSF73 and ecCPSF100 were previously described (32). Cloning of the remaining constructs was performed with ligation independent cloning and plasmids were verified by DNA sequencing. Mutagenesis used PCR with primers containing the desired mutation. **Supplementary Table S1** details all PCR primers used during cloning and mutagenesis, and **Supplementary Table S2** contains the full list of plasmids generated for this study.

### Protein expression

*E. coli* BL21(DE3) *pLysY* (New England Biolabs) were transformed with the various plasmids, and transformant colonies were used for a small-scale overnight culture growth at 37 °C in lysogeny broth (LB) or in terrific broth (TB) supplemented with the corresponding antibiotics. Bacteria from the overnight cultures were used to start 500 mL cultures in LB or TB for natural abundance protein, or in M9 minimal medium supplemented with 1 g/L ^15^NH_4_Cl and 2 g/L ^13^C-glucose. Growth of the 500 mL cultures at 37 °C was followed by induction at an OD_600nm_ of 0.6 with 0.25 mM isopropyl β-D-1-thiogalactopyranoside (IPTG), and protein expression continued for 16 h at 25 °C. For amino acid-specific labelling of isoleucine, valine or leucine, 100 mg/L [^13^C,^15^N]-Ile, [^13^C,^15^N]-Val, or ^13^C,^15^N-Leu was added to the M9 medium 30 minutes prior to induction. Specific labelling of deuterated amino acids used either 100 mg/L [^2^H]-Phe in 500 mL M9, or a combination of 100 mg/L [^2^H]-Tyr and 100 mg/L [^2^H]-Trp in a culture size of 250 mL. Cells were harvested by centrifugation at 4500 x *g* for 20 min at 4^0^C, resuspended in lysis buffer containing 5 mM imidazole, 50 mM Tris (pH 7.5), 500 mM NaCl, 5 % (v/v) glycerol, and stored at -80 °C in the presence of added lysozyme.

### Protein purification

After thawing on ice, resuspended cell pellets were sonicated and soluble protein was separated from cellular debris by centrifugation at 20000 x *g* for 30 min at 4 °C. The supernatant was filtered through a GD/X 0.7 μm filter (GE Healthcare Life Sciences) and loaded onto 2 mL Nuvia IMAC Ni-charged resin (Bio-Rad Laboratories). The resin was washed with 10 column volumes of buffer containing 5 mM imidazole, 50 mM Tris (pH 7.5), 500 mM NaCl, 5 % (v/v) glycerol followed by 5 volumes of the same buffer but with 25 mM imidazole. Protein elution used the same buffer with 500 mM imidazole. Fractions containing the tagged domain were pooled and exchanged to the initial buffer containing 5 mM imidazole by using a PD10 desalting column (GE Healthcare Life Sciences). His-tagged TEV protease (0.1 mg/ml final concentration) was added for overnight cleavage at 4 °C. The protease, hexahistidine tag and any uncleaved protein was removed by a second passage through the Nuvia IMAC Ni-charged resin. The purified samples were concentrated with Vivaspin centrifugal concentrators with cut-off corresponding to the considered fragments (Merck Millipore Corporation) to a volume of 500 μL and exchanged with a NAP-5 column (GE Healthcare Life Sciences) to a buffer consisting of 20 mM Tris (pH 7.5), 150 mM NaCl, 2 mM DTT and concentrated to the desired volume. Protein concentrations were determined by absorbance at 280 nm with extinction coefficients obtained using ProtParam (http://web.expasy.org/protparam).

### Limited proteolysis

Following co-expression of ecCPSF73 and ecCPSF100, histidine-tagged complex was purified on Nuvia IMAC resin. 100 μg of eluted complex were incubated with Trypsin at 0.001 mg/ml final concentration in wash buffer (5 mM imidazole, 50 mM Tris (pH 7.5), 500 mM NaCl, 5 % (v/v) glycerol). Aliquots were collected every 10 min and the reaction was stopped by adding SDS-PAGE loading blue. After 60 min, the samples were loaded on 15% SDS-PAGE for analysis. In a second set of experiments, the reactions were stopped after the optimal incubation time to maximise the generation of the stable fragments and samples were sent to the mass spectrometry platform for molecular mass and peptide identifications.

### NMR spectroscopy

NMR spectra were recorded at 298 K using a Bruker Neo Avance spectrometer at 700 MHz or 800 MHz, equipped with a standard triple resonance gradient probe or cryoprobe, respectively. Bruker TopSpin version 4.0 (Bruker BioSpin) was used to collect data. NMR data were processed with NMR Pipe/Draw (33) and analysed with Sparky 3 (T.D. Goddard and D.G. Kneller, University of California).

### Chemical shift assignment

We have previously reported the backbone and sidechain assignment for ec73-CTD12/ec100-CTD12 (32) which is available from the Biological Magnetic Resonance Data Bank (http://bmrb.wisc.edu/) under BMRB accession number 51624.

For ec73-CTD3, backbone ^1^H^N^, ^1^H^α^, ^13^C^α^, ^13^C^β^, ^13^C’ and ^15^N^H^ chemical shifts were assigned from a sample of 800 µM [^13^C,^15^N]ec73-CTD3 using 2D ^1^H,^15^N-HSQC, 3D HNCO, 3D HNCACO, 3D HNCA, 3D HNCACB, 3D CBCACONH, 3D HNHA and 3D HACACONH spectra. Aliphatic side chain protons were assigned based on 2D ^1^H,^13^C-HSQC, 3D H(C)(CO)NH-TOCSY (60 ms mixing time), 3D (H)C(CO)NH-TOCSY (60 ms mixing time), and 3D (H)CCH-TOCSY spectra. Assignment of sidechain asparagine δ2 amides and glutamine ε2 amides used a 3D ^15^N-HSQC-NOESY (120 ms mixing time). Aromatic ^1^H chemical shifts were assigned using an 800 uM sample of unlabelled ecCPSF73-CTD3 in 99 % D_2_O with 2D TOCSY (60 ms mixing time) and 2D DQF-COSY spectra.

### Structure calculation

Structure ensembles were calculated using Aria 2.3/CNS1.2 (34) (35). Complete refinement statistics for both structures are presented in **Table 1**.

**Table 1.** NMR and refinement statistics.

|  | ec73-CTD12/<br>ec100-CTD12 | ec73-CTD3 |
| --- | --- | --- |
| **Structures in ensemble** | 20 | 20 |
| **Distance restraints** |  |  |
| Total | 6494 | 1693 |
| Intraresidue | 1453 | 555 |
| Interresidue |  |  |
| Sequential ($\vert i-j\vert =1$) | 975 | 245 |
| Short range ($1<\vert i-j\vert <4$) | 471 | 115 |
| Medium range ($3<\vert i-j\vert <6$) | 232 | 73 |
| Long range ($\vert i-j\vert >5$) | 897 | 173 |
| Intermolecular | 305 | - |
| Ambiguous<sup>a</sup> | 1925 | 472 |
| Hydrogen bond | 236 | 60 |
| **Dihedral angle restraints** |  |  |
| Backbone ($\phi, \psi$) | 403 | 106 |
| Sidechain ($\chi_1, \chi_2$) | 127 | 43 |
| **Residual dipolar coupling restraints** |  |  |
| RDC | 165 | - |
| **Structure statistics, mean ± SD** |  |  |
| **Restraint violations** |  |  |
| Distance ($\text{\AA}$) | $0.033 \pm 0.009$ | $0.013 \pm 0.000$ |
| Dihedral angle ($^\circ$) | $0.80 \pm 0.05$ | $0.94 \pm 0.07$ |
| RDC $Q^b$ | $0.17 \pm 0.01$ | |
| **Deviations from idealized geometry** |  |  |
| Bond lengths ($\text{\AA}$) | $0.004 \pm 0.000$ | $0.003 \pm 0.000$ |
| Bond angles ($^\circ$) | $0.72 \pm 0.06$ | $0.45 \pm 0.01$ |
| Impropers ($^\circ$) | $2.8 \pm 0.3$ | $1.2 \pm 0.1$ |
| **Ramachandran plot (%)<sup>c</sup>** |  |  |
| Most favored | 95.6 (91.5) | 93.1 (83.2) |
| Additionally favored | 4.4 (7.4) | 5.6 (14.3) |
| Generally allowed | 0.0 (0.8) | 0.8 (1.4) |
| Disallowed | 0.0 (0.3) | 0.5 (1.1) |
| **rmsd to average structure ($\text{\AA}$)<sup>c</sup>** |  |  |
| Backbone ordered | $0.8 \pm 0.2$ | $0.4 \pm 0.2$ |
| Heavy atom ordered | $1.2 \pm 0.2$ | $0.8 \pm 0.2$ |

<sup>a</sup> Distance restraints in ARIA from the sum of two or more overlapping NOESY crosspeaks
<sup>a</sup> Calculated as described in (56)
<sup>c</sup> Determined using the PSVS validation suite (57) (58) Values are reported for ordered residues, with the Ramachandran analysis for all residues included in parentheses.

For the heterodimer formed by ec73-CTD12/ec100-CTD12, ^1^H distances were first obtained using NOE crosspeaks from 3D ^15^N-HSQC-NOESY (120 ms mixing time) and 3D aliphatic ^13^C-HSQC-NOESY (120 ms mixing time) spectra using sample protein concentrations of 880 and 230 µM, respectively. For aromatic ^1^H, the majority of distance restraints were obtained from a 458 μM unlabelled sample in 99 % D_2_O using a 2D ^1^H,^1^H-NOESY (120 ms mixing time) spectrum. Similar to the strategy used in the original chemical shift assignment (32), manual NOE crosspeak assignment was assisted by comparison to a 2D ^1^H,^1^H-NOESY spectrum collected on a sample of ^2^H-Phe (150 μM), and a spectrum from a sample including both ^2^H-Tyr and ^2^H-Trp (85 μM). Additional ^1^H distances were obtained by using a 3D aliphatic ^13^C-HSQC-NOESY (120 ms mixing time) spectrum collected from a 625 µM sample in which only the isoleucine was ^13^C-labelled, as well as a spectrum from a 694 µM sample in which both valine and isoleucine were ^13^C-labelled. Starting at iteration four, residual dipolar coupling (RDC) values were included based on interleaved spin state-selective TROSY spectra from a sample of 230 µM [^2^H,^15^N]ec73-CTD12/ec100-CTD12, without and with the addition of 18 mg/ml Pf1 phage. RDC-based intervector projection angle restraints relate to D_a_ and R values of 10 and 0.64, respectively. Protein dihedral angles were obtained by using TALOS-N (36) and SideR (37)(38). Hydrogen bond restraints (two per hydrogen bond) were introduced following an initial structural calculation and include only protected amides in the centre of secondary structure elements. Final ensembles were refined in explicit water and consisted of the 20 lowest energy structures from a total of 250 calculated models.

For CTD3 of ecCPSF73, the ^1^H distances were obtained using NOE crosspeaks from a sample of 800 µM [^13^C,^15^N]ec73-CTD3 using a 3D ^15^N-HSQC-NOESY (120 ms mixing time) spectrum, and on the same sample in 99 % D_2_O using a 3D aliphatic ^13^C-HSQC-NOESY (120 ms mixing time). Distance restraints, in particular for the aromatic ^1^H, were also obtained from an 800 μM sample of unlabelled ec73-CTD3 in 99 % D_2_O from a 2D ^1^H,^1^H-NOESY (120 ms mixing time) spectrum. Protein dihedral angles were obtained by using TALOS-N (36) and SideR (37)(38). Hydrogen bond restraints (two per hydrogen bond) were introduced following an initial structural calculation and include only protected amides in the centre of secondary structure elements. Final ensembles were refined in explicit water and consisted of the 20 lowest energy structures from a total of 100 calculated models.

### Pull-down assays

All proteins were purified as described above with the exception that the tagged proteins were used directly after the first Ni-NTA purification without elution. Protein concentrations were determined by Coomassie staining. Equal amounts of individual tagged proteins were used for each experiment. Pull-down assays were performed by mixing tagged proteins on Ni-NTA beads with excess untagged constructs in binding buffer (20 mM sodium phosphate, pH 7.4, 150 mM NaCl, 2 mM ß-mercaptoethanol) and incubated for 1 h at 20 °C. Beads were then washed 3 times with the same buffer, and bound proteins were eluted and analyzed on 18 % SDS-PAGE.

### Model predictions by AlphaFold2

Predicted structural models were generated with AlphaFold2 (39) as implemented in the ColabFold server (40). For each complex, five models were generated using default parameters, although with a maximum number of recycles set at 12. Of the five structures, the one with the highest pTMscore was chosen as the representative model. The per-residue pLDDT scores for each representative model are shown in **Supplementary Figures S2** and **S6**. The sequences chosen for protein prediction (including Uniprot ID, amino acid range) are: hsCPSF73 (Q9UKF6, 460-684), hsCPSF100 (Q9P2I0, 651-782), hsSympk (Q92797, 539-1100); scYsh1 (Q06224, 474-779), scCft2 (Q12102, 718-859), scPta1 (Q01329, 559-785), scSyc1 (Q08553, 1-188); spYsh1 (O13794, 464-757), spCft2 (O74740, 637-797), spPta1 (Q10222, 445-670); tcCPSF73 (Q6BCB3, 482-762), tcCPSF100 (Q6BCB1, 626-802), tcSympk (A0A7J6XPR2, 573-928); ceCPSF73(Q95PY8, 458-707), ceCPSF100 (O17403, 633-843), ceSympk (O16929, 525-1143); atCPSF73 (Q9C952, 465-693), atCPSF100 (Q9LKF9, 602-739), atSympk (Q9SFZ8, 474-1092); dmCPSF73 (Q9VE51, 464-684), dmCPSF100 (Q9V3D6, 608-756), dmSympk (Q8MSU4, 516-1101); ecCPSF73 (Q8SUE4, 452-643 and 568-643), ecCPSF100 (Q8SRZ4, 525-639), ecSympk (Q8SVB3, 161-385). Full prediction details, along with all structural models, are accessible at: https://www.nmrbordeaux.org/Data/Thore_et_al_AlphaFold_models.zip.

### Electrophoretic mobility shift assays

The binding of DNA and RNA to CPSF protein constructs used a Tris-glycine gel system (36) and non-specific nucleotide sequences as used in a previous study (41). Protein samples were prepared in 20 mM Tris (pH 7.5), 150 mM potassium acetate, 1 mM EDTA and 10% glycerol, with bromophenol blue added to aid in gel loading. 3’Cy3 labelling was incorporated into the forward strands of the DNA (AGGGTCTCCATTTTGAAGCATGC-Cy3) and RNA (AGGGUCUCCAUUUUGAAGCAUGC-Cy3) during chemical synthesis on an Expedite 8909 (PerSeptive Biosystems). Double-stranded oligonucleotides were prepared with unlabelled reverse strands (GCATGCTTCAAAATGGAGACCCT and GCAUGCUUCAAAAUGGAGACCCU) by heating 50 μM mixed samples to 98 °C, and allowing them to slowly cool to room temperature. For each assay, 500 nM of the Cy3-labelled single- or double-stranded oligonucleotide was used without or with 50 μM protein, and analyzed after incubation for 30 minutes. During the incubation step, polyacrylamide gels were prepared with a final concentration of 10 % acrylamide:bisacrylamide (37.5:1), 300 mM Tris– HCl (pH 8.8), 0.1% (w/v) ammonium persulfate and 0.1% (v/v) TEMED. The wells were washed with water after a polymerization of 10 min, and the gel was pre-run at room temperature for 30 min at 100 V in a running buffer containing 27 mM Tris, 192 mM glycine, 1 mM EDTA and pH 8.3. Following a second cleaning of the wells, 2 μl of each sample was loaded and the gel was run at 100 V for an additional 30 min at room temperature. Visualization of Cy3 fluorescence used a Typhoon Trio+ imager (GE Healthcare) with a 580 nm filter, 532 nm laser, normal sensitivity, photomultiplier tube setting of 450 V and 100 micron resolution. Preparation of images used ImageQuant TL v8.1.0.0 with default visualization parameters. In a final step, the gels were stained with Coomassie blue to visualize the protein.

## RESULTS

### C-terminal heterodimer of CPSF73 and CPSF100 has two independent modules

To understand the molecular basis of heterodimer formation by the C-termini of CPSF73 and CPSF100, we initially focused on the human protein complex. Although protein production was observed, significant instability and rapid precipitation complicated their study. A search for possible alternative species identified excellent protein production and stability for the complex formed from the orthologous proteins of the parasite *Encephalitozoon cuniculi*. The initial complex was made by co-expression in *E. coli* with the full C-terminal region of both ecCPSF73 (residues 452-643, hereafter ec73-CTD123; **Figure 1A**) and ecCPSF100 (residues 525-639, hereafter ec100-CTD12). Characterization of this heterodimer by NMR spectroscopy reveals a well-folded complex with dispersed peaks in the ^1^H,^15^N-HSQC spectrum (**Supplementary Figure S1A**). A notable property of the spectrum is a subset of dispersed peaks with higher intensity, which suggests the presence of a smaller folded module independent of the larger complex. To identify the protein sequences corresponding to the two modules we used limited trypsin proteolysis on the heterodimer sample with an N-terminal His6-tag on ec73-CTD123. Subsequent Ni-column purification showed that an N-terminal fragment of His6-ec73-CTD123 retained binding to a mostly intact ec100-CTD12, and mass spectrometry identifies an ecCPSF73 trypsin cleavage site at Lys571 (**Supplementary Figure S1B**). We therefore made two new constructs of ecCPSF73 based on secondary structure prediction and the trypsin digest results (**Figure 1A**): ecCPSF73 from residue 452-567 (ec73-CTD12), and another from 567-643 (ec73-CTD3). Characterization by NMR spectroscopy of the new heterodimer prepared by co-expression of ec73-CTD12 and ec100-CTD12 showed that a stable and folded complex was produced (**Figure 1B**). In addition, the isolated CTD3 from ecCPSF73 also produced a stable and folded protein by NMR spectroscopy (**Figure 1C**). Using these two samples we began structure determination to access their atomic details.

**Figure 1.**
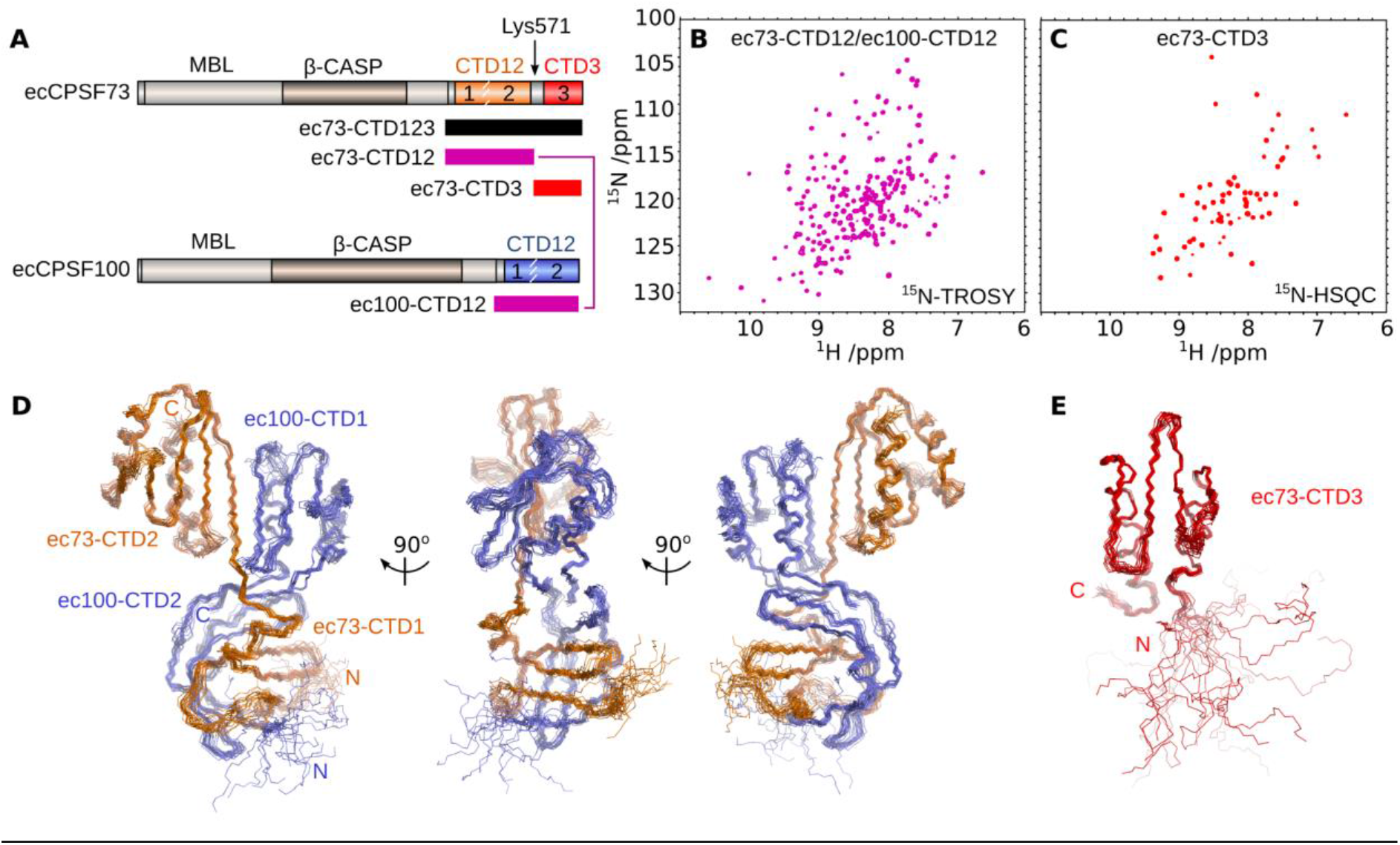
The C-terminal heterodimer of CPSF73/CPSF100 is formed from two independent modules. (**A**) Domain architecture of ecCPSF73 and ecCPSF100. Construct boundaries are shown below, and the location of trypsin-sensitive ecCPSF73 residue Lys571 is also indicated. (**B**) ^1^H,^15^N-TROSY spectrum of [^15^N]-labelled ec73-CTD12/ec100-CTD12. (**C**) ^1^H,^15^N-HSQC spectrum of [^15^N]-labelled ec73-CTD3. (**D**) Backbone line representation of the ensemble of 20 structures calculated for the CTD12-CTD12 heterodimer, with ecCPSF73 in orange and ecCPSF100 in violet. (**E**) Ensemble of 20 structures calculated for CTD3 from ecCPSF73.

### Solution structures of the two C-terminal modules

To determine the structure of ec73-CTD12/ec100-CTD12 we initiated a crystallization screen, as well as preliminary data collection by NMR spectroscopy. Due to an absence of crystals, we prioritized determination of the solution structure by using NMR spectroscopy. Despite the high quality of the NMR spectra based on ec73-CTD12/ec100-CTD12 (**Figure 1B**), its overall size at 27 kDa required additional NMR approaches to simplify and reduce ambiguity in the chemical shift assignment. A usual strategy would involve isotopic-labelling of only one protein at a time in the complex, however attempts to produce the heterodimer from separated bacterial expressions were unsuccessful. We previously took advantage of amino acid-selective isotopic labelling of the complex, with samples [^13^C,^15^N]-labelled on Ile, Val or Leu to help with backbone and side chain assignments of these residues (32). It was also apparent early on that aromatic residues would define the hydrophobic core of this complex, and thus to ensure unambiguous assignment we used samples with ring-deuterated Phe or Tyr/Trp to selectively remove cross-peaks in the spectra. Near-complete chemical shift assignment was therefore obtained for the entire complex (32). Continuing this selective-labelling approach, we obtained a large number of NOESY-derived distance restraints, as well as orientational restraints from residual dipolar couplings in a phage-aligned sample (**Table 1**). These data were complemented by dihedral angle predictions to generate an initial ensemble of structures by using ARIA2.3/CNS1.2 (34, 35). Inspection of these ensembles and observation of amide ^1^H^N^ NOESY cross-peaks enabled us to include additional hydrogen bond restraints in the final structure calculation. The resulting ensemble of 20 structures was obtained with a consistent overall architecture and good statistics (**Figure 1D**; **Table 1**).

The smaller C-terminal module only consists of the CTD3 from ecCPSF73, and in this case the NMR spectroscopy and structure calculation used a standard approach to derive the ensemble of 20 structures (**Figure 1E**; **Table 1**).

### Overview of the CTD12-CTD12 heterodimer

As can be seen in the ensemble of structures, the entire CTD12 region from both ecCPSF73 and ecCPSF100 forms a single folded complex (**Figure 1D**). A notable feature of the complex is the extensive contacts made between secondary structure elements of ec73-CTD12 and ec100-CTD12 (**Figure 2A,B**). The buried interface is calculated to be 1910 Å^2^ using the PISA server (42), and the high degree of intermolecular contacts between the two proteins creates an overall architectural stability as evident by global fit of the RDC values (**Table 1**). Despite this single folded structure, it is possible to consider three sub-domains within the complex (**Figure 2A,B**): the CTD1-CTD1 β-barrel, a central region, and the CTD2-CTD2 segment.

**Figure 2.**
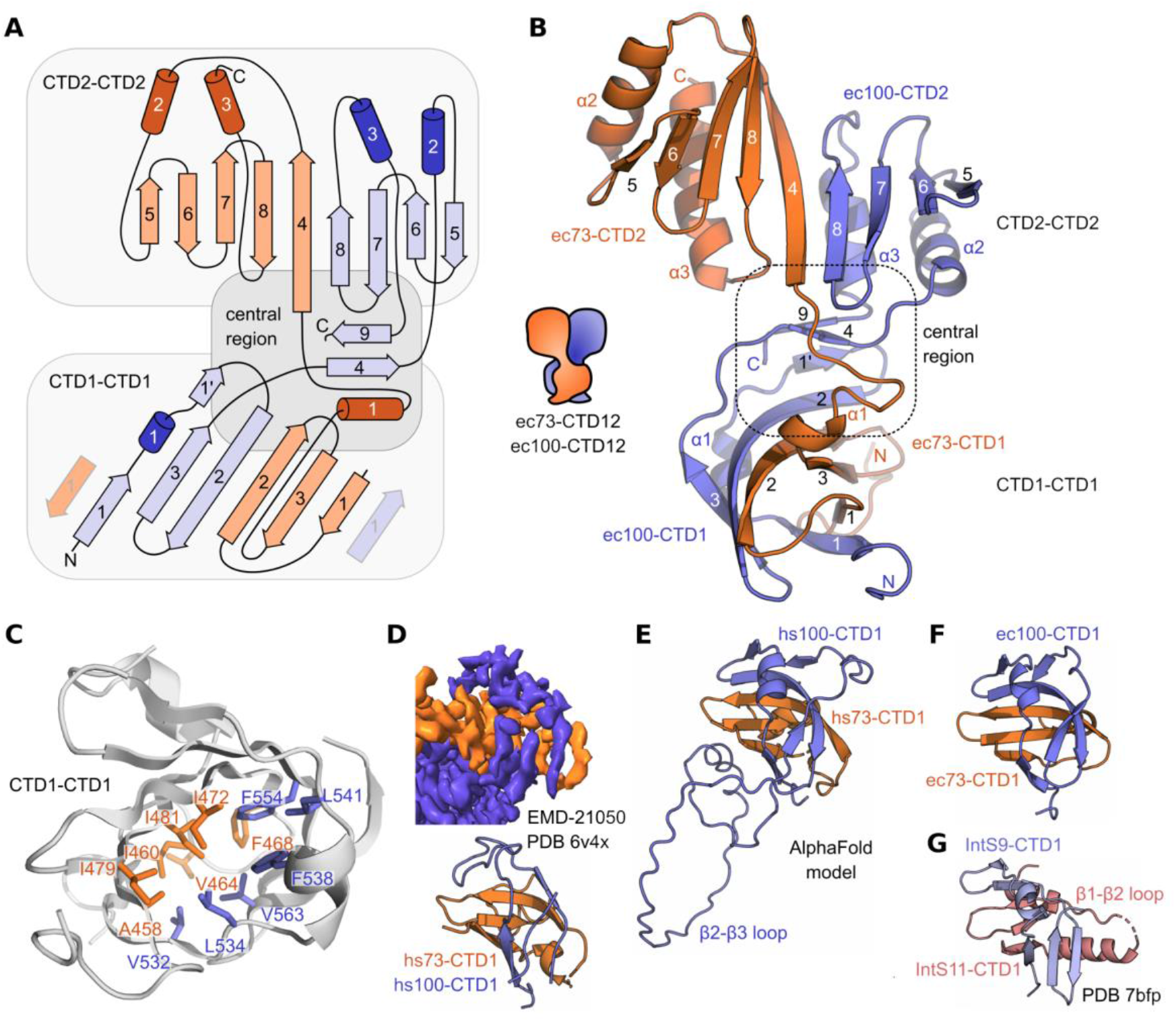
Structural details of the CTD12-CTD12 heterodimer. (**A**) Schematic representation with annotated α-helices (cylinders) and β-strands (arrows) coloured as in Figure 1D, with the three sub-regions indicated. (**B**) Representative model from the NMR ensemble, with annotated secondary structure elements. (**C**) Hydrophobic residues in the core of the CTD1-CTD1 β-barrel shown as sticks and annotated. The view is rotated 180° around the vertical axis as compared to (B). (**D**) CTD1-CTD1 β-barrel from the human histone pre-mRNA 3’ processing complex from EMD-21050, and the derived PDB 6×4v (22). (**E**) CTD1-CTD1 β-barrel from the AlphaFold2 model (**Supplementary Figure S2A**). (**F**) *E. cuniculi* CTD1-CTD1 β-barrel with the same orientation as panels D, E and G. (**G**) CTD1-CTD1 β-barrel from human Integrator (PDB ID 7bfp)(26).

### CTD1-CTD1 β-barrel

Three β-strands from each CTD1 region of ecCPSF73 and ecCPSF100 come together to form the CTD1-CTD1 β-barrel structure (**Figure 2A,B**). Specifically, the β2 strands from each protein form an extended antiparallel arrangement, followed by intramolecular antiparallel β2-β3 contacts and parallel β3-β1 contacts. The barrel is closed by the antiparallel interaction of the two β1 strands, with the β1 strand from ecCPSF100 interrupted by a short helical segment. The hydrophobic residues in the core are shown in **Figure 2C**. For the human complex, the CTD1-CTD1 β-barrel was observed at low resolution in the cryo-EM study of the histone pre-mRNA 3’-end processing machinery (**Figure 2D**) (22), however the limited resolution prevented a detailed comparison. To this end, we have used AlphaFold2 (39) to expand the atomic model of the entire human CPSF73/CPSF100 C-terminal complex (**Supplementary Figure 2A,B**). The top-ranked model displays excellent agreement with the cryo-EM map for the CTD12-CTD12 region (**Supplementary Figure S2C**). Using this model, we observe that the human CTD1-CTD1 β-barrel is highly similar to the *E. cuniculi* NMR structure (**Figure 2E,F**). To determine the extent of similarity with other species, we then predicted models for the CPSF73/CPSF100 C-terminal complex from four diverse model organisms (*Trypanosoma cruzi*, *Caenorhabditis elegans*, *Arabidopsis thaliana* and *Drosophila melanogaster*) as well as the corresponding Ysh1/Cft2 complex from the yeast *Saccharomyces cerevisiae* and *Schizosaccharomyces pombe* (**Supplementary Figure S2D**). As expected, an overall structural similarity is clear from these additional models. However, it is also evident that large inserts are present at differing locations within the β-barrels (**Supplementary Figure S2E**), such as the large β2-β3 loop insert from hs100-CTD1 (**Figure 2E**). As a final comparison, cryo-EM data are available for the IntS11 and InstS9 proteins within the human Integrator complex, such as shown for PDB ID 7bfp (**Figure 2G**)(26). A similar architecture is clearly observed despite the limited resolution, but with a helix inserted within the IntS11-CTD1 β1-β2 loop.

### CTD2 belongs to the TBP domain family

The CTD2-CTD2 region of CPSF73/CPSF100 had already been proposed to form in a similar manner to the C-terminal heterodimer of IntS11/IntS9 (23) and suggested by the lower resolution cryo-EM data (22). Consistent with this observation, we find in our structure that the CTD2-CTD2 region forms an extended β-sheet formed by β-strands β4-β8 of ecCPSF73 and β5-β8 of ecCPSF100, with four α-helices on the opposing side (**Figure 2A,B**). To identify key residues, we created a sequence alignment for the C-terminal regions (**Supplementary Figure 3A,B**). The alignment required use of our predicted models with the PROMOLS3D structure-based protocol (43), to account for the variable loops in the sequences. Highest conservation is observed for residues at the interface between the two CTD2 domain (**Figure 3A,B**), and indicate an importance for ionic interactions between the absolutely conserved Asp555 of ecCPSF73, and Arg629 of ecCPSF100 that is an Arg in all sequences except for a Lys in scCft2. Surrounding residues are also conserved. A similar ionic bridge appears to be key for the Integrator proteins (**Figure 3C**) as shown by the interaction of Glu581 from IntS11 and Arg648 of IntS9 (23).

**Figure 3.**
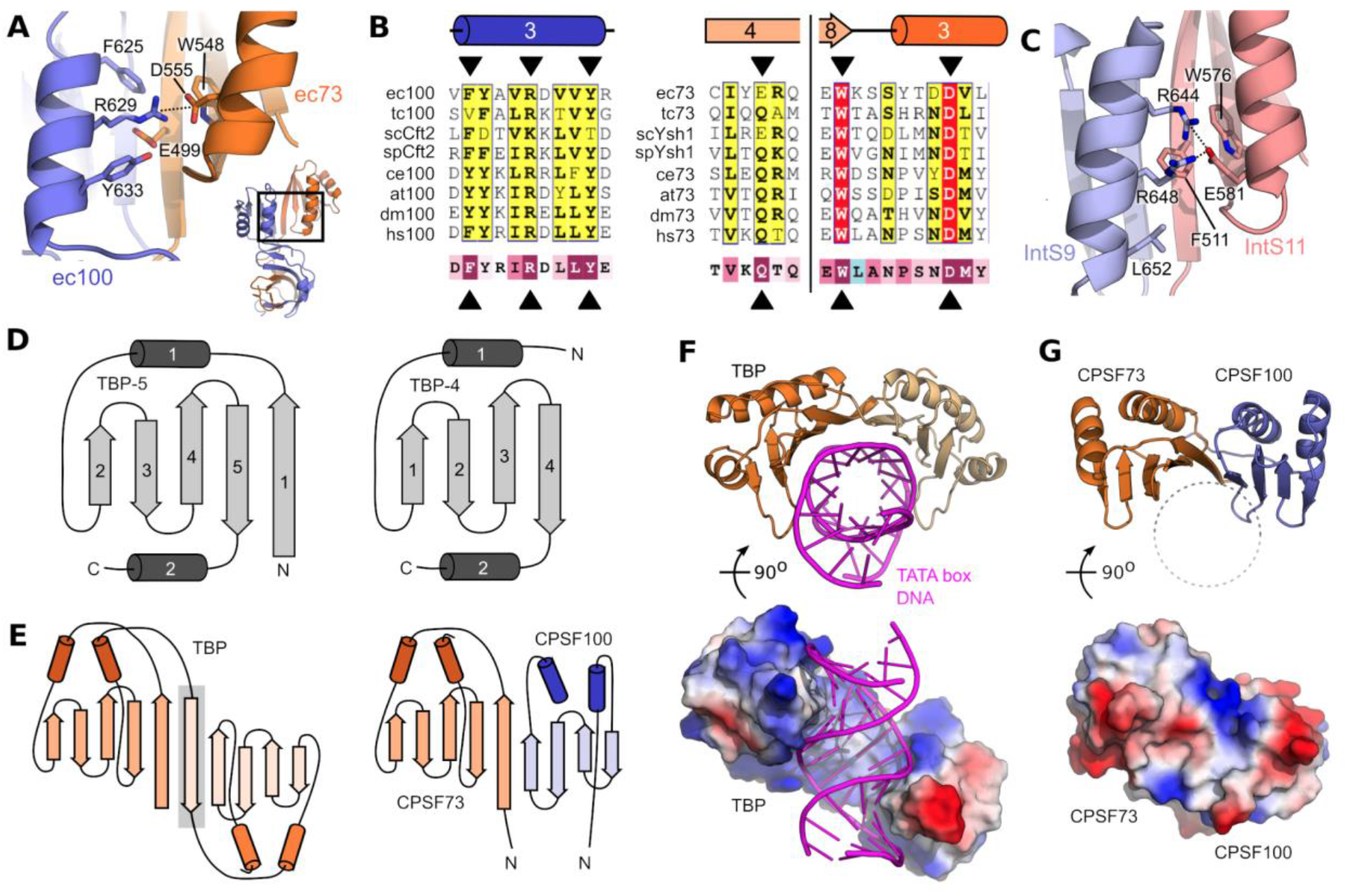
The CTD2 domains are part of the TBP domain family. (**A**) Close up view of the interdomain contacts between ec73-CTD2 and ec100-CTD2. (**B**) Conservation of residues at the CTD2-CTD2 interface (indicated by black triangles) using PROMOLS3D (43) and ESPRIPT (59). The bottom line shows conservation of each residue based on the human sequences and generated by ConSurf (46). Colour spectrum goes from poorly conserved (teal) to highly conserved (magenta). Full alignments and complete details are found in **Supplementary Figure S3.** (C) Close up view of the interdomain contacts between hsIntS11-CTD2 and hsIntS9-CTD2 (PDB ID 5v8w) (23). (D) Five-stranded (left) and four-stranded (right) TBP domain folds. (**E**) The TBP protein is composed of tandem five-stranded TBP domains, as compared to the five-stranded TBP domain from CPSF73 and the four-stranded TBP domain from CPSF100. The additional β-strand in TBP is highlighted in grey. (**F)** Cartoon representation (top) and surface charge representation (bottom) of the TBP-DNA complex (PDB ID 1cdw) (48). (**G**) The corresponding cartoon and surface charge view for ec73-CTD2/ec100-CTD2.

In order to identify additional structural homologs of the CTD2-CTD2 region and the entire CTD12-CTD12 heterodimer, we used the FoldSeek webserver (44)(https://search.foldseek.com/) since it is well-suited for multi-domain structures, and takes into account both the PDB as well as the AlphaFold Protein Structure Database. As expected, we identified the IntS9/IntS11 C-terminal complex from the Integrator cryo-EM structure (FoldSeek E-value of 0.07 for PDB 7bfq; (26)), as well as predicted models for CPSF73 or CPSF100 from numerous species. However, we also found structural similarity between the CTD2-CTD2 segment and the TATA-box binding protein (TBP), with E-values comparable to that of Integrator. The TBP domain family shares a core architecture of α-β-β-β-β-α, with or without an additional β-strand at the N-terminus (**Figure 3D**). The CTD2 domain thus fits into this family, and the overall CTD2-CTD2 architecture is remarkably similar to TBP except that the N-terminal β-strand is missing in CPSF100 (**Figure 3E**). In addition to TBP, the domain has been found in bacterial DNA glycosidases and the archeal ribonuclease H3, and may have evolved from a distant single domain protein ancestor (45). Each of these proteins are involved to different degrees in nucleic-acid binding: TBP has high specificity for the DNA TATA box (**Figure 3F**), the domain from ribonuclease H3 is non-sequence specific for DNA-RNA duplexes, and DNA glycosidases use a separate domain to bind the DNA target (**Supplementary Figure 4A**). Unlike TBP, the presence of several acidic residues centred in the β-strands would argue against a nucleic acid-binding function for the CTD2-CTD2 β-sheet in CPSF73/100 (**Figure 3G**). Indeed, using NMR spectroscopy and a previously established electrophoretic mobility shift assay protocol (41), we did not detect any binding by the CTD12-CTD12 heterodimer to single-stranded or double-stranded DNA and RNA ligands (**Supplementary Figure 4D,E**).

### Hydrophobic contacts in the central region

In our high-resolution structure of the CTD12-CTD12 heterodimer, we were able to observe all residues within the complex, and therefore we were able to detect a large hydrophobic core that joins the CTD1-CTD1 and CTD2-CTD2 regions (**Figure 4A**). In the context of the complete CTD12-CTD12 heterodimer (**Figure 2A,B**), these residues connect the upper surface of the CTD1-CTD1 β-barrel, ecCPSF73 linker helix α1, ecCPSF100 linker strand β4, bottom of the CTD2-CTD2 segment, and the final β-strand (β9) of ecCPSF100. This central hydrophobic core is not unique to *E. cuniculi*, as seen in the high conservation of corresponding residues mapped onto the human CTD12-CTD12 model (**Figure 4B**). The final residue of ecCPSF100 is particularly interesting in this context, as the C-terminus position is strictly conserved in length and is invariably a hydrophobic residue (**Supplementary Figure 3B).** This hydrophobic C-terminal side chain (Ile639 in ecCPSF73, **Figure 4A**) is structured in the CTD12-CTD12 ensemble, and directly contributes to the central hydrophobic core. The AlphaFold models suggest that a similar situation exists for other organisms, such as for Val782 at the C-terminus of hsCPSF100 (**Figure 4B**). Re-analysis of the initial crystal structure of the IntS11-IntS9 CTD2-CTD2 heterodimer (PDB 5v8w)(23) shows that intermolecular contacts already exist outside of the primary CTD2-CTD2 interface. For example, the final residue of hsIntS9 (Phe658) is visible in the crystal structure, and is surrounded by neighbouring hydrophobic residues N-terminal to the CTD2 domains (**Figure 4C**). Despite the absence of residues from the top of the CTD1-CTD1 β-barrel, this partial central core is robust enough to form in a manner similar to that observed in the complete ec73-CTD12/ec100-CTD12 heterodimer.

**Figure 4.**
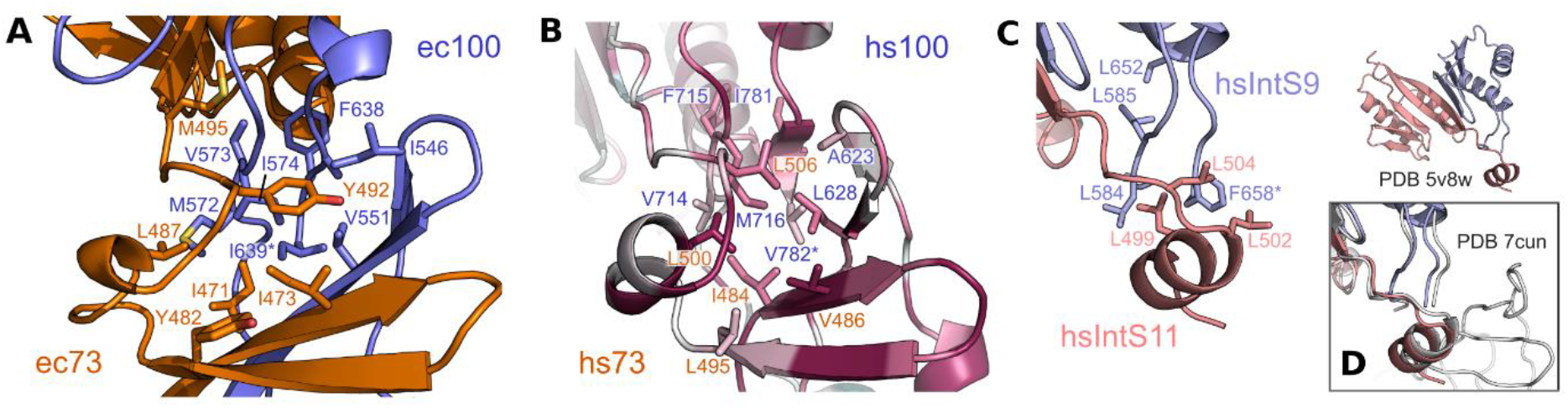
Hydrophobic core of the CTD12-CTD12 central region. (**A**) Closeup view of the central region with residue sidechains annotated and shown in stick representation. The C-terminal residue is indicated with an asterisk (**B**) Central region from an AlphaFold model of the human CTD12-CTD12 complex. Annotated hydrophobic sidechains are coloured based on their ConSurf conservation score (most conserved in magenta, average in white, and least conserved in teal). See **Supplementary Figure 2** for complete ConSurf analysis. (**C**) Annotated sidechains from the partial central region of the isolated CTD2-CTD2 complex from human Integrator proteins IntS11 and IntS9 (PDB iD 5v8w; (23)). (D) For reference, the boxed view shows a superposition onto the intact Integrator proteins within the structure of the human Integrator-PP2A complex (PDB ID 7cun; (47)).

### CTD3 of CPSF73 binds Symplekin

In contrast to the larger CTD12-CTD12 module, the second module of the CPSF73/CPSF100 C-terminal complex is only composed of ecCPSF73 CTD3. A representative model from the ensemble of structures shows that ec73-CTD3 is composed of a single folded domain that is joined by a flexible linker to the rest of ecCPSF73 (**Figure 5A,B**). The α-β-β-β-β-α architecture again places it in the TBP domain family, and this time FoldSeek indicates the closest structural homology to the archeal ribonuclease H3 N-terminal domain (RMSD of 2.5 Å; **Supplementary Figure S4B**). In keeping with a four-stranded TBP domain fold, ec73-CTD3 also displays structural similarity to CTD2 in ecCPSF100 (**Supplementary Figure S4C**), despite sharing only 13 identical residues (∼20% of the sequence) between the two folded domains.

**Figure 5.**
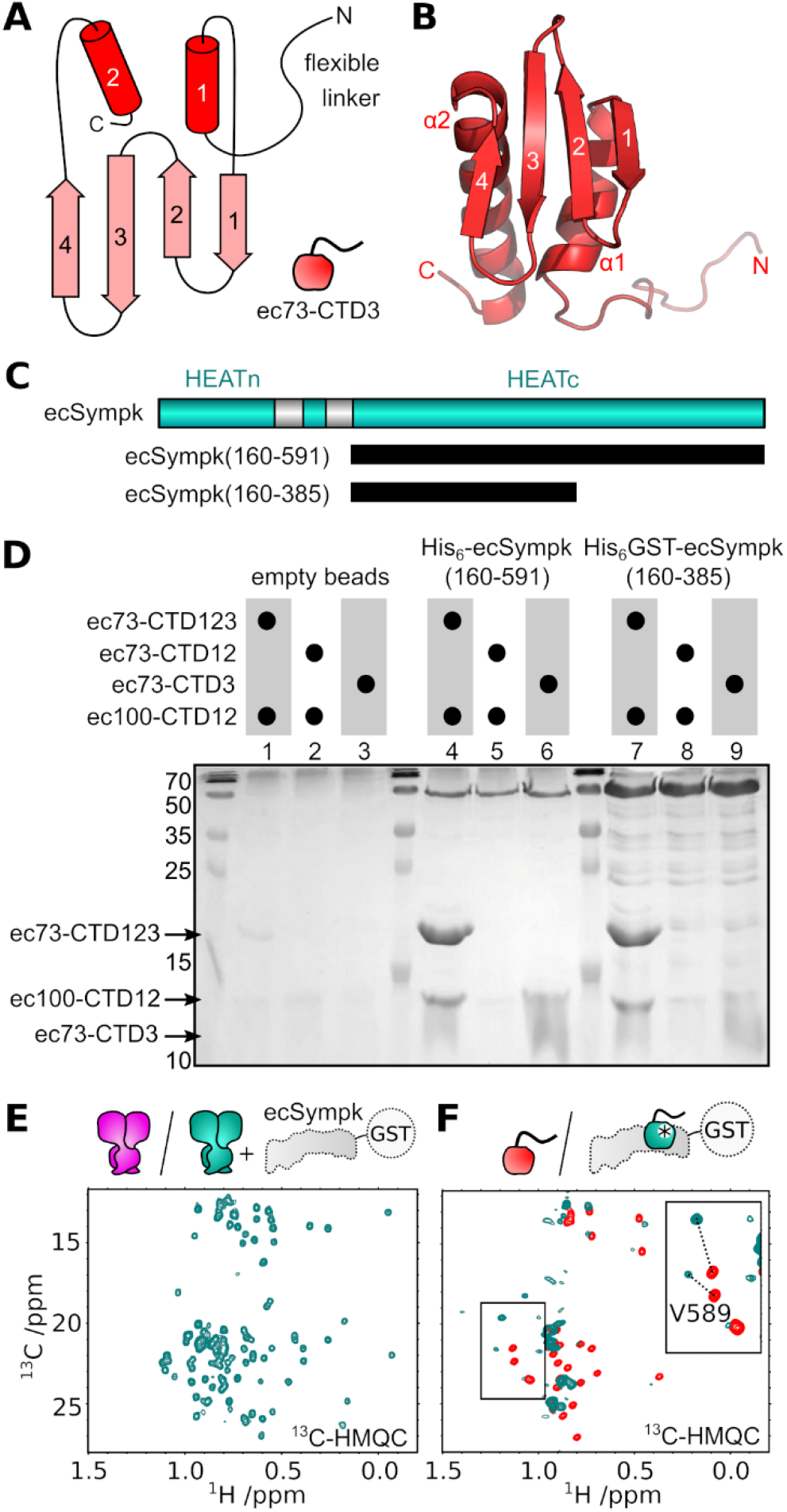
CTD3 of CPSF73 is required to bind Symplekin. (**A**) Schematic of the ec73-CTD3 domain. (**B**) Representative model of ec73-CTD3 with annotated secondary structure elements. (**C**) Domain architecture of ecSympk with construct boundaries. (**D**) SDS-PAGE gel following pull-down of various ec73 and ec100 samples using empty beads, the full HEATc, or a truncated HEATc construct of ecSympk. (**E**) Methyl group region of ^13^C-HMQC spectra for the CTD12-CTD12 heterodimer without (magenta) and with (cyan) two molar equivalents of unlabelled GST-ecSympk(160-385). No changes are detected between the two spectra, supporting an absence of interaction. (**F**) Similar experiments with isolated ec73-CTD3 (red) and upon addition of GST-ecSympk(160-385) (cyan) shows chemical shift perturbation of most methyl groups. The insert shows large perturbation (dotted lines) of the two methyl crosspeaks from Val589.

Similar to the CTD12-CTD12 heterodimer, we failed to detect a general ability of ec73-CTD3 to bind nucleic acids (**Supplementary Figure S4F**). However, it was shown in the cryo-EM structure of the human histone pre-mRNA 3’ processing complex that CPSF73 CTD3, and also the CTD2-CTD2 region, are likely in direct contact with the Symplekin protein (22). We therefore decided to see if our ecCPSF73 and ecCPSF100 constructs (**Figure 1A**) indeed bind to the *E. cuniculi* Symplekin (ecSympk). In the human complex, it is the C-terminal HEAT repeats of Symplekin that are involved in the interaction, and therefore we started with the same region of ecSympk **(Figure 5C**; residues 160-591). Using a pull-down assay, we found that the entire C-terminal complex of ec73-CTD123/ec100-CTD12 was able to interact with ecSympk(160-591) (**Figure 5D, Supplementary Figure S5A**). Furthermore, we found that although the isolated ec73-CTD3 construct retained the ability to bind, there was no detectable interaction with the CTD12-CTD12 heterodimer. We further refined the interacting region in ecSympk to residues 160-385 (**Figure 5D, Supplementary Figure S5B**). To confirm our findings, we used NMR spectroscopy to test for binding to ecSympk(160-385). Once again, we failed to detect any interaction between the CTD12-CTD12 heterodimer and ecSympk (**Figure 5E**), whereas binding by ec73-CTD3 was clearly evident due to the large degree of chemical shift perturbation upon the addition of GST-ecSympk(160-385) (**Figure 5F**).

### Conserved interaction with Symplekin

Given the importance of ec73-CTD3 in the binding of ecSympk(160-385), we chose to generate a model of this complex in order to design mutants to disrupt this interaction. The AlphaFold model of the complex (**Supplementary Figure S6A,B**) indicates that the main interface is hydrophobic and centred on helix α1 of ec73-CTD3 (**Figure 6A**). We chose to introduce the acidic residue Glu in place of Val589 in the middle of this interface, or Leu585 towards the edge. Based on NMR spectroscopy, both mutant proteins were well-folded (**Figure 6B**) but only V589E prevented this interaction. The ability of the ec73-CTD3 V589E mutant to disrupt the interaction with ecSympk was confirmed in a pull-down experiment (lane 11, **Figure 6C**). This assay also shows that Leu585 is only able to disrupt the interface in the context of the helix α1 L585K,N592D double mutation (lanes 10 and 12, **Figure 6C**). The corresponding hydrophobic surface on ecSympk is predicted to include Phe212, Tyr215, Phe216 and Val245 (**Figure 6A**), and indeed a V245D mutation is able to weaken the complex (**Supplementary Figure S5C**).

**Figure 6.**
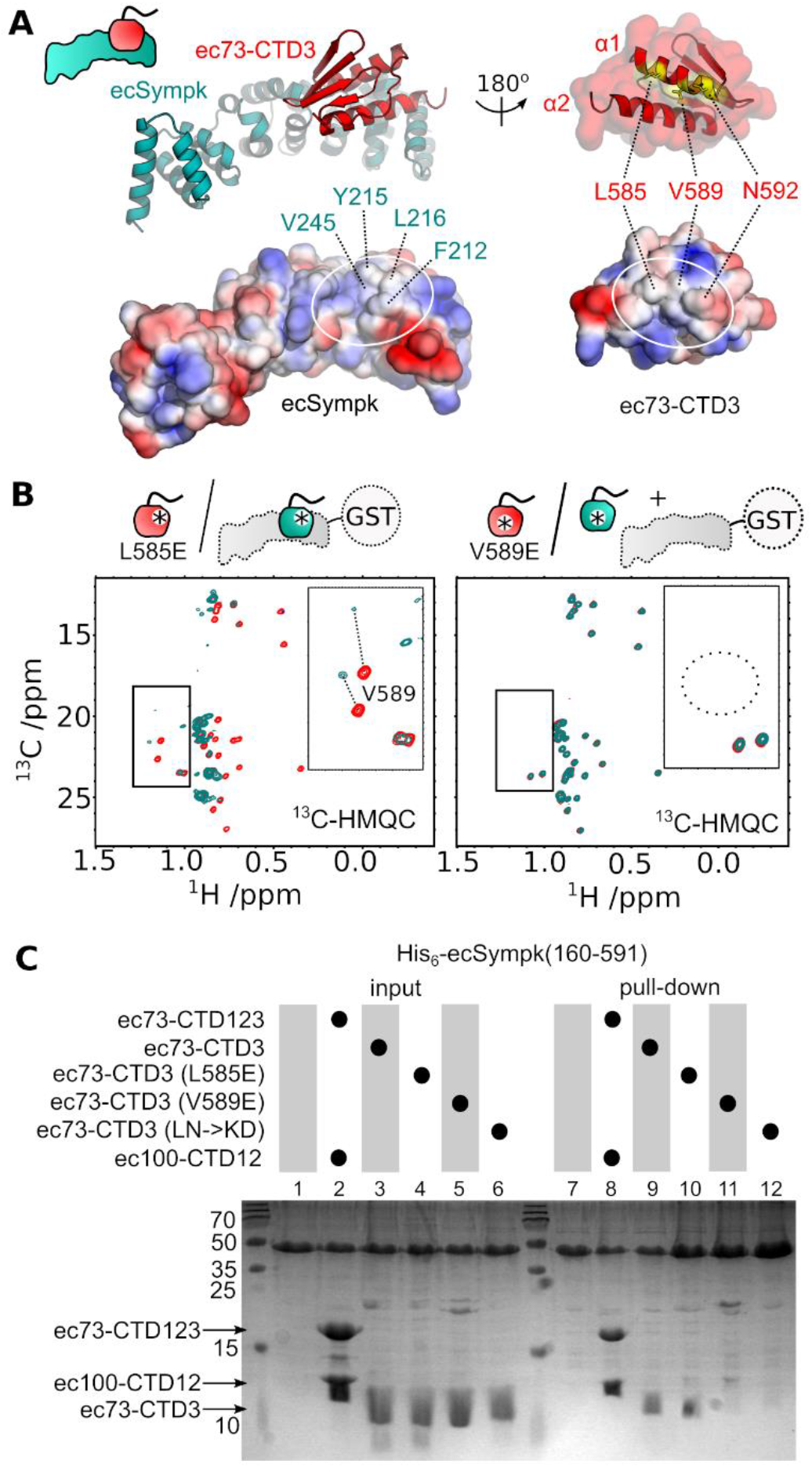
Mutagenesis of the interface between ec73-CTD3 and ecSympk. (**A**) AlphaFold model of the complex between ec73-CTD3 and ecSympk(160-385). (**B**) Similar to Figure 5F, methyl group region in ^13^C-HMQC spectra for ec73-CTD3 mutants L585E (left) and V589E (right), in the absence (red) and presence (cyan) of unlabelled GST-ecSympk(160-385). Both ec73-CTD3 mutant proteins are folded, but only V589E prevents binding to ecSympk. The mutation of Val589 is also confirmed by the disappearance of its two methyl groups in the spectrum (dotted circle) when changed to a glutamate. (**C**) SDS-PAGE gel from a pull-down experiment between ecSymp(160-591) and the wildtype ec73-CTD123/ec100-CTD12 complex, or a series of wild-type and mutant ec73-CTD3.

To explore a wider conservation of the CTD3-Sympk interaction, we expanded our AlphaFold models to include the C-terminal region of human Symplekin in complex with hs73-CTD123 and hs100-CTD12 (**Figure 7A**, **Supplementary Figure S6C**). There is excellent agreement of this model with the cryo-EM map of the histone pre-mRNA 3’-end processing complex (**Supplementary Figure S6D**)(22). Using this model, we were able to look for surface residue conservation by using the ConSurf program (46). The region on Symplekin in contact with hs73-CTD3 is enriched in conserved residues (white circle, **Figure 7B**), and is surrounded by residues that are highly variable. Furthermore, the hs73-CTD3 surface in contact with hsSympk is enriched in conserved hydrophobic residues (right, **Figure 7B**), consistent with the interface we observed for ec73-CTD3. Residue conservation for the CTD2 domains is instead mostly limited to residues that form the CTD2-CTD2 interface (as shown in **Figure 3A,B**). Additional models for the CPSF73-CPSF100-Sympk C-terminal complexes in *T. cruzi*, *C. elegans*, *A. thaliana*, and *D. melanogaster* show a similar arrangement of CPSF73 CTD3 onto Symplekin (**Supplementary Figure S6E**). A conserved interaction for the yeast homologues are also seen from models determined for Ysh1-Cft2-Pta1 C-terminal complexes from *S. cerevisiae* and *S. pombe* (**Supplementary Figure S6E**). Using *S. cerevisiae* as an example, the surface region on scPta1 in contact with scYsh1-CTD3 is clearly enriched in conserved residues (**Figure 7C,D**).

**Figure 7.**
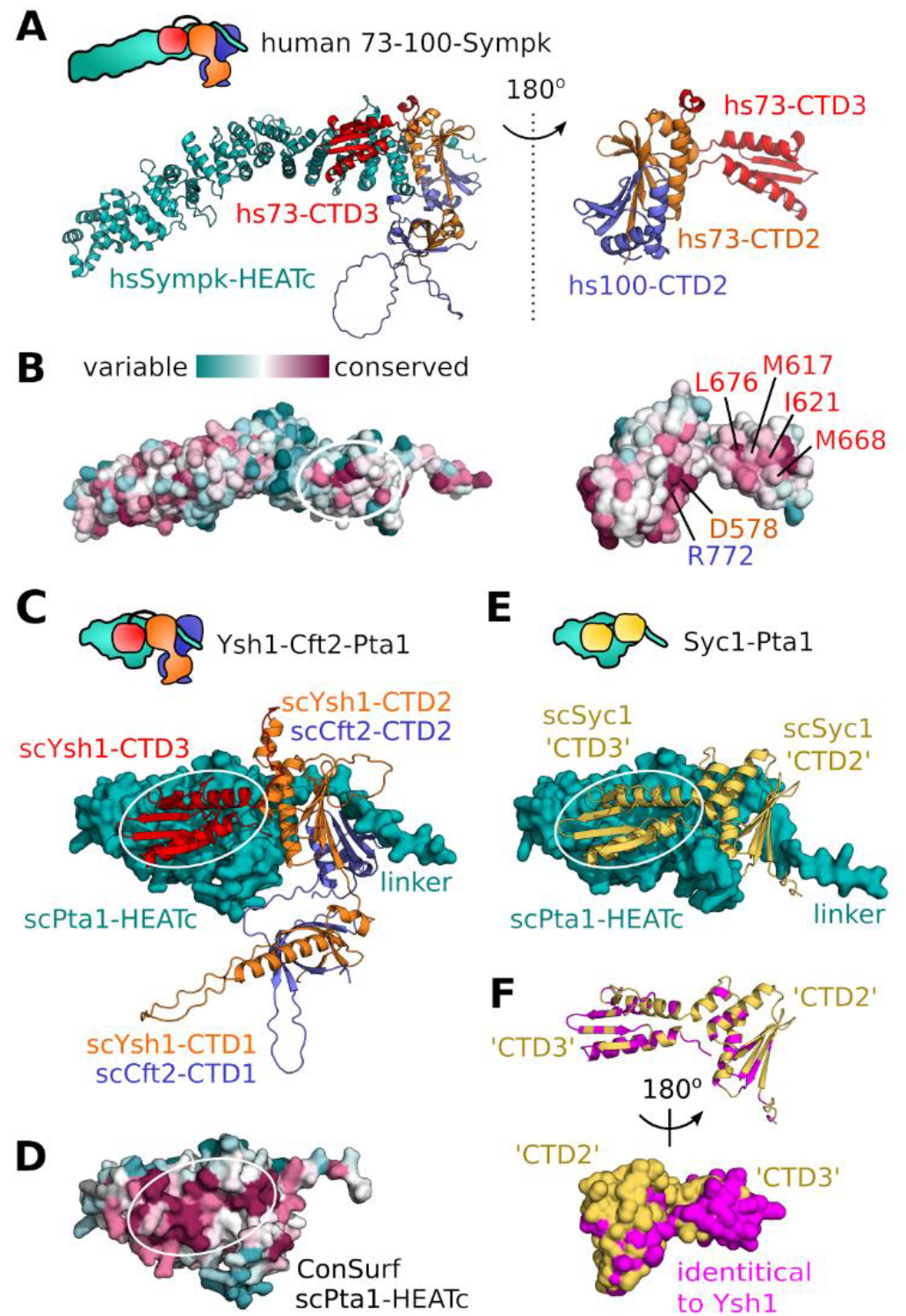
Conserved interaction between CTD3 and Symplekin/Pta1. (A) AlphaFold model of the human CPSF73-CPSF100-Symplekin C-terminal trimer (Supplementary Figure S6C,D). (B) Surface residue conservation using ConSurf (46) with the location of CTD3 binding on Sympk circled (left), and the corresponding surface shown on hs73-CTD3 (right). Surface hydrophobic residues on hsCTD3 are annotated in red. Also annotated are hs73-CTD2 Asp578 and hs100-CTD2 Arg772 at the CTD2-CTD2 interface. (C) Predicted model of the *S. cerevisiae* Ysh1-Cft2-Pta1 C-terminal complex (Supplementary Figure S6E). The C-terminal HEAT repeats of scPta1 is shown in surface representation, and the other proteins are shown as cartoons. (D) Surface residue conservation of scPta1. (E) Predicted model of the *S. cerevisiae* Syc1-Pta1 complex (Supplementary Figure S6F). (F) Identical residues between *S. cerevisiae* Syc1 and Ysh1 (Supplementary Figure 6G) are coloured in magenta. All residues in the ‘CTD3’ domain of Syc1 that contact Pta1 are identical with scYsh1.

## DISCUSSION

We have determined atomic details for the two structured modules formed by the C-termini of CPSF73 and CPSF100. The largest module comprises a heterodimer of the CTD1 and CTD2 from each protein, with extensive intermolecular contacts creating the 27 kDa complex. For full structure determination by NMR spectroscopy, a complex of this size remains challenging, particularly without an option to selectively label only one of the peptides. We instead took advantage of several amino-acid type labelling strategies to simplify the NMR data analysis, reduce ambiguity in the crowded spectra, and obtain a large number of structural restraints (as shown in **Table 1**). Using NMR spectroscopy on the isolated CTD1-CTD2 heterodimer allowed us to access full details of the intra- and inter-molecular contacts, and to detect aspects of protein dynamics. Apart from high-resolution details of the CTD1-CTD1 and CTD2-CTD2 regions of the CPSF73-CPSF100 complex, we observed a significant number of hydrophobic residue contacts between these components to form an extended hydrophobic core. The entire CTD1-CTD2 heterodimer therefore behaves as a single structural unit, which likely persists within the assembled CPSF.

Focussing on the β-barrel formed by the CTD1 of both proteins, we confirm structural similarity to the partial models built for this region from cryo-EM data of the histone pre-mRNA 3’ processing machinery (22), as well as IntS11 and IntS9 within Integrator (26)(47)(29). Based on both predicted models and alignments from diverse model organisms (**Supplementary Figures S2** and **S3**), the β-barrel region typically includes loop insertions of varying length, such as the extensive 62 amino-acid loop between strands β2 and β3 in human CPSF100. The function of these species-specific loops is currently unknown. We note, however, that the CTD1-CTD2 regions from *E. cuniculi* CPSF73 and CPSF100 lack these extended loops, which may have been a factor in their favourable *in vitro* behaviour.

Moving to the CTD2 region, we find that the CTD2-CTD2 dimer is indeed similar to the heterodimer structure first determined for the isolated CTD2 of Integrator proteins IntS11 and IntS9 (23). A search for additional structure similarity shows that the CTD2 falls into the larger family of TATA-box binding protein (TBP) domains (45). The CTD3 of CPSF73 is also a TBP domain, with a four-stranded architecture such as seen in the CTD2 of CPSF100. The TBP domain family is typically associated with nucleic-acid binding proteins, with direct contact to DNA observed for the tandem TBP domains of TATA-box binding protein (48, 49), and the binding of an RNA-DNA duplex by the TBP domain of archeal ribonuclease HIII (50). In both cases, the contact to nucleic acids is by the β-sheet surface. In contrast, we did not find any detectable binding to single- or double-stranded oligonucleotides for the CTD2 of CPSF73-CPSF100 or indeed for CPSF73 CTD3 (**Supplementary Figure S4**). Although it is possible that binding is restricted to a specific motif, the fact that there are few conserved basic residues on the β-sheet surfaces argues against a nucleic acid-binding function. A similar case is observed for IntS11 and IntS9, for which there is non-specific ssRNA binding detected along one side of the CTD1-CTD1 β-barrel within the IntS4-IntS9-IntS11 trimer, but no interaction with the CTD2 domains (26). Instead, the AlphaFold2 predicted models show that the peptide region of Symplekin/Pta1 that connects the N- and C-terminal domains tends to lie across the centre of the CTD2-CTD2 β-sheet surface (**Figure 7A,B**; **Supplementary Figure S6**). It may be that the TBP domain in these cases have a preference for binding peptides, and thus may help in the overall architecture of the CPSF and histone mRNA 3’ processing machinery.

In contrast to the larger CTD12-CTD12 heterodimer, the second structured module in the CPSF73-CPSF100 C-termini is just the single CTD3 domain of CPSF73. In this case, there is a clear role for the TBP domain in protein-protein interaction. However, this interaction with Symplekin involves the two α-helices on the opposite side of CTD3 from the β-sheet surface. Although we did not detect a high affinity interaction between the Symplekin CTD and the CPSF73-CPSF100 heterodimer module, it is again the α-helices of the CTD2 domains that make contact with Symplekin in the assembly of the three proteins. Our NMR data provided atomic details of the CPSF73 CTD3 and also allowed us to probe the specific interaction to Symplekin. A hydrophobic surface centred on Val589 in the first α-helix of ec73-CTD3 (corresponding to Met617 in human CPSF73, **Figure 7A**) is key to the interaction with Symplekin, and a mutation to Glu preserves the fold but abolishes the interaction. For the corresponding surface on Symplekin, we combined sequence analysis (ConSurf) and predicted models (AlphaFold2) to identify a consistent patch of conservation where CPSF73 CTD3 appears to bind (**Figure 7A**).

The key role of CTD3 in the interaction between CPSF73 and Symplekin also marks an important difference in comparison to Integrator. For IntS11 (a paralogue to CPSF73) there is no CTD3 domain, and therefore this mode of interaction is not possible. Indeed, except for a common role in the assembly architecture, Symplekin and the Integrator protein IntS4 share little sequence similarity and only resemble each other by containing a series of HEAT repeats (25). Consistent with this difference, the main contact between IntS11-IntS9 and the C-terminus of IntS4 is on the opposite side of the CTD12-CTD12 heterodimer, as compared to Symplekin and CPSF73-CPSF100 (26). Further specification of the CPSF73-CPSF100 interaction with Symplekin lies in the fact that despite some sequence conservation, CPSF73 can not form a heterodimer with IntS9 (27). Altogether, the intimate and specific dimerization of CPSF73 and CPSF100, via the CTD12-CTD12 heterodimer, ensures that only Symplekin is able to interact in the formation of the trimeric CCC/mCF core cleavage module.

The available cryo-EM data for human CPSF and the histone pre-mRNA 3’ processing machinery, along with several studies of Integrator, have allowed us to interpret our findings on CPSF73, CPSF100 and Symplekin in various contexts. In contrast, the orthologous regions in the yeast proteins, namely the C-termini of Ysh1, Cst2 and Pta1, do not have reported structural data. Nevertheless, our analysis of sequence conservation and predicted models provide confidence that similar interactions of the three proteins are preserved within the yeast cleavage and polyadenylation factor (CPF). A similarity between the yeast and human proteins has been previously validated by the ability of some mammalian factors to replace yeast proteins (51)(10). In CPF, Ysh1 and Cft2 belong to the nuclease module within the core CPF (CPF_core_) (30) and their C-terminal regions likely form a CTD12-CTD12 heterodimer as observed for CPSF73 and CPSF100 (**Figure 7C**). The model of the Ysh1-Cft2-Pta1 ternary complex supports a shared mode of interaction to Pta1 driven by the CTD3 domain of Ysh1, with further contacts to the CTD2 α-helices of Ysh1 and Cft2. This interaction helps link the CPF nuclease module to the phosphatase module. As seen for Symplekin, there is minimal direct contact of Pta1 with Cft2 in the predicted model, which in yeast may have an additional consequence. The APT complex (named for Associated with PTa1) is a separate 3’ processing machinery in *S. cerevisiae* that displays a preference for sn/snoRNA (52). The Syc1 protein is key to forming the APT complex, and appears to bind Pta1 in a mutually exclusive manner to Ysh1 (53). The two domains of full-length Syc1 are most closely related to CTD2 and CTD3 of Ysh1/CPSF73 (54). An AlphaFold model of the Syc1-Pta1 complex shows a remarkable similarity in the position and mode of Pta1 binding between Syc1 and scYsh1-CTD23 (**Figure 7E**, **Supplementary Figure 6F**). This model would explain how the single Syc1 protein binds to Pta1 in a mutually exclusive manner to Ysh1 (and Cft2), to prevent inclusion of the nuclease and polymerase modules in the APT complex. The model again highlights the important role played by the CTD3 domain. The second domain in Syc1, which we name ‘CTD3’ due to similarity to scYsh1-CTD3, has complete conservation of all residues that interact with Pta1 (**Figure 7F**).

The structural details we have determined for the C-terminal complex of CPSF73 and CPSF100, as well as the trimer formation with Symplekin, fills a previous gap in the atomic description of CPSF in general, and CCC/mCF in particular. Our high-resolution data allows for a detailed description of the complete CTD12-CTD12 heterodimer, as well as the CPSF73 CTD3 structure and interaction with Symplekin. The next steps will use this information to understand the role of these interactions in the function of the CCC/mCF, taking advantage of structure-guided mutations. The consequence of post-translational modifications will add an additional layer of regulation to be addressed. Related to the current study, sumoylation has been observed for human CPSF73 on Lys462 and Lys465 which we place along the first β-strand of CTD1 (55). This modification would clearly impact assembly into CPSF or CCC/mCF. Sumoylation also occurs on Lys535, located in the loop between the first and second β-strands of CTD2, which may have additional implications. These and other protein modifications may be important in certain developmental or disease states. Finally, further study of the complexes from different organisms, including our use of the parasite *E. cuniculi*, may discover functionally important species-specific aspects within the CCC/mCF such as in infectious parasites. Such differences could in turn be exploited to derive new therapeutic avenues to treat infections and disease.

## DATA AVAILABILITY

The structure ensemble of the ec73-CTD12/ec100-CTD12 complex has been deposited at the Protein Data Bank (http://www.ebi.ac.uk/pdbe/) with accession ID 8BA1, and the ensemble of ec73-CTD3 structures with accession ID 8B7T. Corresponding chemical shift assignments of ec73-CTD3 have been deposited in the Biological Magnetic Resonance Data Bank (http://bmrb.wisc.edu/) under BMRB accession number 34760. Prediction details and structural models are accessible at: https://www.nmrbordeaux.org/Data/Thore_et_al_AlphaFold_models.zip.

## AUTHOR CONTRIBUTIONS

Molecular biology and protein production were performed by all authors. CDM collected and processed the NMR data and performed the structure calculations. ST and CDM analysed the NMR data. ST performed the pull-down experiments. ST, SF and CDM wrote the manuscript and prepared the figures. All authors gave final approval for the submission and agreed to be held accountable for the work.

## ACKNOWLEDGEMENTS

We thank Axelle Grélard, Estelle Morvan, and the structural biology platform at the Institut Europèen de Chimie et Biologie for access to the NMR spectrometers, equipment, and technical assistance, and Post Sai Reddy for collecting the mass spectrometry data. We also thank Lori Passmore and colleagues for discussion.

