## Supplementary Data for "Molecular details of the CPSF73-CPSF100 C-terminal heterodimer and interaction with Symplekin"

#### SUPPLEMENTARY TABLES

**Supplementary Table S1.** List of PCR primers.

| Function | Direction | Primer sequence (5'→3') |
|---|---|---|
| Clone ec73-CTD123 from genomic DNA | Forward | gaaggagatatacatATGCTGGCAAAGGACAACGAG |
| | Reverse | tctagactattaggatccCTAATCTCTCAAATATAT |
| Clone ec73-CTD12 from ec73-CTD123 | Forward | CTTTACTTCCAGGGCTCCCTGGCAAAGGACAACGAG |
| | Reverse | TCTAGACTATTAGGATCCCTAGTCAGATATCATCTTGC |
| Clone ec100-CTD12 from genomic DNA | Forward | tattttcagggctccatgTCTGATGTTTCGATGGG |
| | Reverse | tctagactattaggatccTCATATAAAGGCCGAGCTTCT |
| Clone ec73-CTD3 from ec73-CTD123 | Forward | ctttacttccagggctccGGGTTGAAGACCAAGAGT |
| | Reverse | tctagactattaggatccCTAATCTCTCAAATATAT |
| Clone ecSympk (160-591) from genomic DNA | Forward | ctttacttccagggccatATGCCTGTCAGCGAGTCTTTC |
| | Reverse | tctagactattaggatccCTATTTATTACCAAAC |
| Clone ecSympk (160-385) from genomic DNA | Forward | ctttacttccagggccatATGCCTGTCAGCGAGTCTTTC |
| | Reverse | tctagactattaggatccTTACCGGACCGCCCTTCTTTT |
| Clone ecSympk (385-591) from genomic DNA | Forward | ctttacttccagggccatATGGGCCAGCTCCGGGTCTAT |
| | Reverse | tctagactattaggatccCTATTTATTACCAAAC |
| ec73-L585E mutation | Forward | GGAAGAAGGAGGAATTGGTGAGTGTTTTGAAG |
| | Reverse | CAAAACACTCACCAATTCCTCCTTCTTTCCCATC |
| ec73-V589E mutation | Forward | CTATTGGTGAGTGAGTTGAAGAACCACTTCATGG |
| | Reverse | GGATAGTGGACGAGATAAATGAGTGTATCAAAAAG |
| ec73-I639E mutation | Reverse | CTCCTAGGGCTAGCTCTAGACTAATCTCTCAAATATTCCG CCTCTATCTTTTTGATAC |
| ec73-L585K, N592D mutations | Forward | AAGAAGGAGaaATTGGTGAGTGTTTTGAAGgACCACTTCA TG |
| | Reverse | CATGAAGTGGTcCTTCAAAACACTCACCAATttCTCCTTC TT |

**Supplementary Table S2.** Reagents and resources.

| REAGENT or RESOURCE | SOURCE | IDENTIFIER |
|---|---|---|
| Bacterial and Virus Strains | | |
| 5-alpha Competent E. coli | New England Biolabs | Cat# C2987H |
| T7 Express lysY Competent E. coli | New England Biolabs | Cat# C3010I |
| BL21 Rosetta2 Competent E. coli | Sigma-Aldrich | Cat#71397 |
| Chemicals, Peptides, and Recombinant Proteins | | |
| Nuvia™ IMAC Resin | BioRad | Cat# 7800800 |
| TEV protease | In house produced | N/A |
| Deposited Data | | |
| ec73-CTD12/ec100-CTD12 chemical shifts | BioMagResBank (BMRB) | 51624 |
| ec73-CTD12/ec100-CTD12 structural ensemble | Protein Data Bank (PDB) | 8BA1 |
| ec73-CTD3 chemical shifts | BioMagResBank (BMRB) | 34760 |
| ec73-CTD3 structural ensemble | Protein Data Bank (PDB) | 8B7T |
| Oligonucleotides | | |
| PCR oligos | Sigma/Eurofins | Supplementary Table S1 |
| DNA, RNA ligands | In house produced | Supplementary Fig S4 |
| Recombinant DNA | | |
| pET-MCN | Novagen | N/A |
| pET-MCN-GST | Novagen | N/A |
| pET-CDF | Novagen | N/A |
| ecCPSF73 | genomic DNA | Based on genome sequence from (60) |
| ecCPSF100 | genomic DNA | |
| ecSymplekin | genomic DNA | |
| pET-MCN:ec73(452-643) [ec73-CTD123] | This paper | N/A |
| pET-MCN:ec73(452-567) [ec73-CTD12] | This paper | N/A |
| pET-CDF:ec100(525-639) [ec100-CTD12] | This paper | N/A |
| pET-MCN:ec73(567-643) [ec73-CTD3] | This paper | N/A |
| pET-MCN:ecSympk(160-591) | This paper | N/A |
| pET-MCN-GST:ecSympk(160-385) | This paper | N/A |
| pET-MCN:ecSympk(385-591) | This paper | N/A |
| pET-MCN:ec73(567-643)L585E | This paper | N/A |
| pET-MCN:ec73(567-643)V589E | This paper | N/A |
| pET-MCN:ec73(567-643)I639E | This paper | N/A |
| pET-MCN:ec73(567-643)L585K, N592D | This paper | N/A |
| Software and Algorithms | | |
| Topspin | Bruker BioSpin | Version 4.0 |
| NMRPipe | (33) | Version 8.6 |
| Sparky | T.D. Goddard and D.G. Kneller, Univ. of California | Version 3.1.1.3, and NMRFAM-SPARKY 1.470 |
| ARIA | (34) | Version 2.3 |
| CNS | (35) | Version 1.2 |
| PROMOLS3D | (43) | online server |
| ConSurf | (46) | online server |
| Pymol | Schrodinger, LLC | Version 1.4.1 |
| ChimeraX | (61) | Version 1.1 |

### SUPPLEMENTARY FIGURES

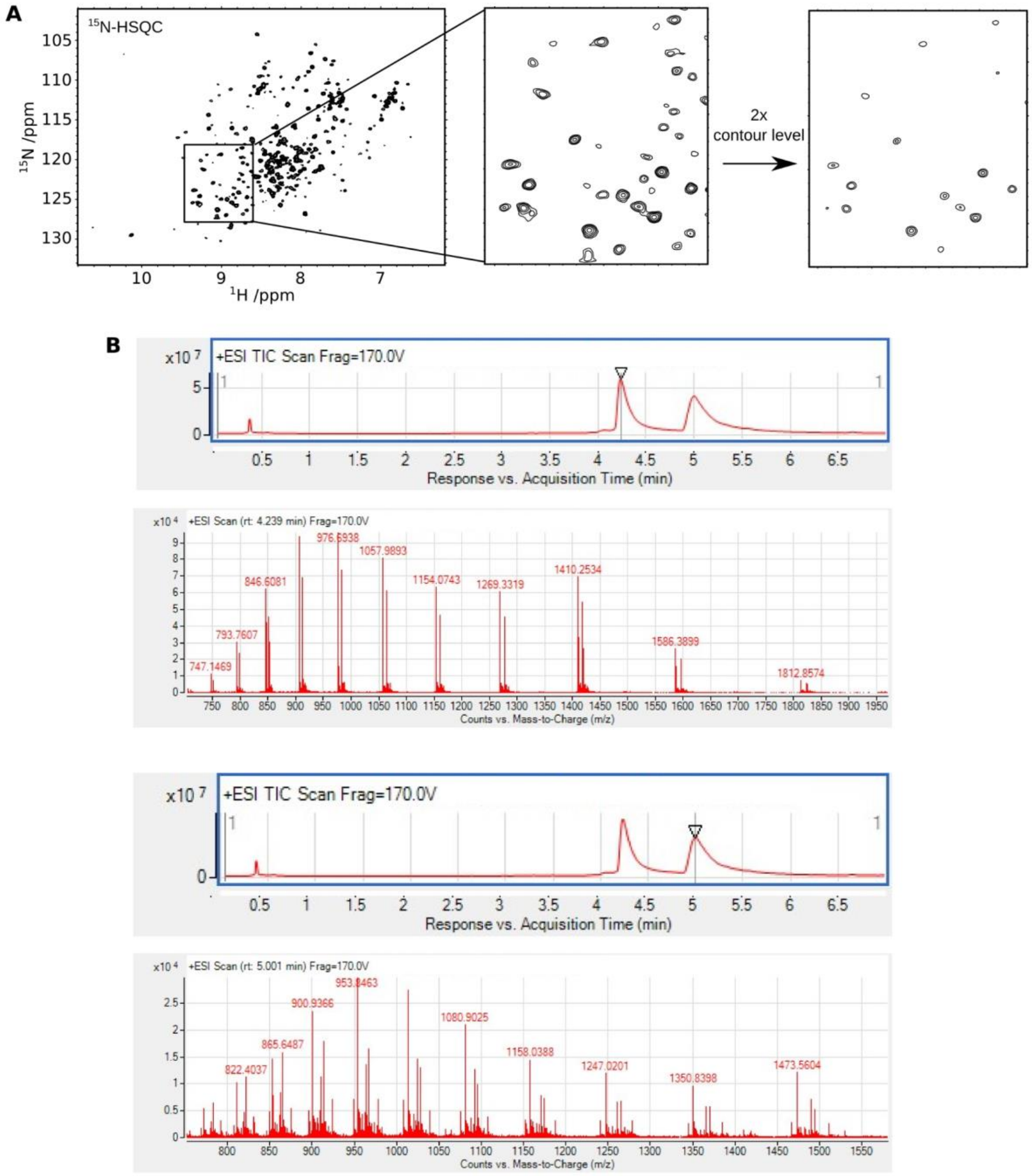


**Supplementary Figure S1**. Definition of the two structural modules in the ec73-CTD123/ec100-CTD12 complex. (**A**) $^{15}$N-HSQC spectrum of [15N]ec73-CTD123/ec100-CTD12, including a close up view of the selected region plotted at the same contour level, and then at twice the threshold. A subset of the peaks have a higher signal-to-noise behaviour. The spectrum was collected at 298 K at a field strength of 700 MHz. (**B**) Mass spectrometry analysis of the complex remaining after limited trypsin proteolysis. (Top) The eluted peptide at 4.24 min has a predominant mass at 11290 Da, corresponding to the complete C-terminus of ecCPSF100 starting with residue Gly537. (Bottom) The peptide eluting at 5.00 min has a predominant mass of 16213 Da, corresponding to the His-tag up until Lys571 of ecCPSF73. Only the region corresponding to CTD1-CTD2 is therefore present in the trypsin-resistant complex.

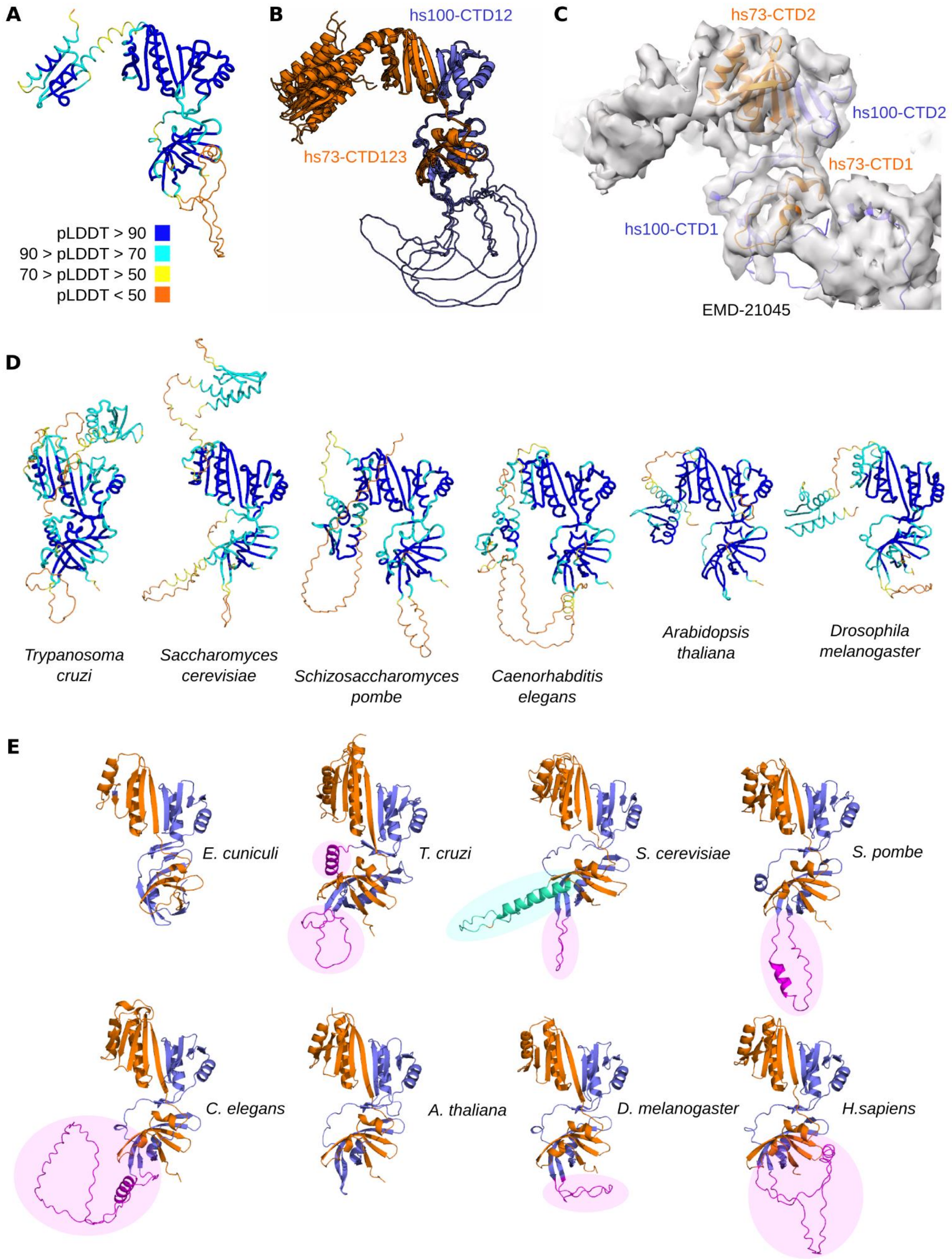


**Supplementary Figure S2**. AlphaFold2 models of the human CPSF73/CPSF100 C-terminal heterodimer and additional model organisms. (**A**) Top pTMscore-ranked AlphaFold2 model of the human CPSF73-CTD123/CPSF100-CTD12 complex, shown in tubes coloured by pLDDT. (**B**) All five predicted models of the human CPSF73-CTD123/CPSF100-CTD12 complex, shown in cartoon and coloured by chain. The overlay highlights uncertainty in the relative position of CPSF73-CTD3, and in the large CPSF100-CTD1 loop. (**C**) AlphaFold2 model of the human CTD12-CTD12 complex (B) docked into the cryo-EM map of the histone pre-mRNA 3'-end processing complex (EMD-21045)(Sun et al., 2020). (**D**)Top pTMscore-ranked AlphaFold2 models of the CPSF73-CTD123/CPSF100-CTD12 complexes from additional model organisms, shown in tubes coloured by pLDDT. (**E**) Cartoon representations of the NMR structures of the CTD12-CTD12 complex from *E. cuniculi*, and taken from the seven AlphaFold2 models. Large loop inserts in CTD1 from various CPSF100/Cst2 are shown in magenta, and the loop insert from *S. cerevisiae* Ysh1 CTD1 in cyan.

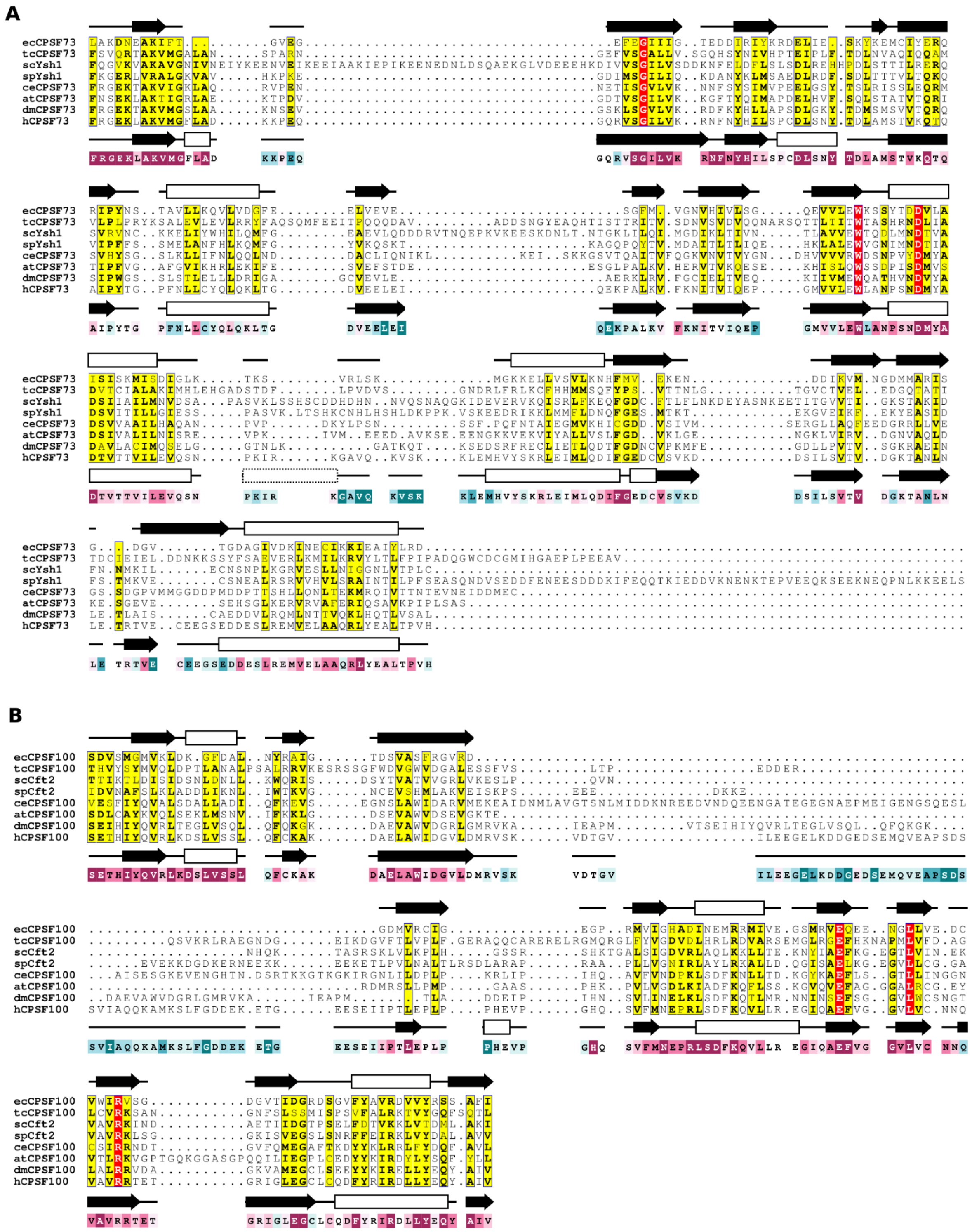

**Supplementary Figure S3**. C-terminal regions of CPSF73 and CPSF100 from *E cuniculi* and model organisms. Sequence alignments for the C-terminal regions of (**A**) CPSF73, (**B**) CPSF100. The alignments were generated by using PROMOLS3D (43) and the AlphaFold2 models from **Supplementary Figure 2**, then formatted by using ESPRIPT (59). The secondary structure elements from the NMR structure of ec73-CTD123/ec100-CTD12 and ec73-CTD3 are shown above the alignment, and those from the AlphaFold2 model of human CPSF73/ CPSF100 are shown below. The final line shows the conservation of each residue based on the human sequences and generated by ConSurf (46). Colour spectrum goes from poorly conserved (teal) to highly conserved (magenta). Species: ec, *Encephalitozoon cuniculi*; tc, *Trypanosoma cruzi*; sc, *Saccharomyces cerevisiae*; sp, *Schizosaccharomyces pombe*; ce, *Caenorhabditis elegans*; at, *Arabidopsis thaliana*; dm, *Drosophila melanogaster*; h, human.

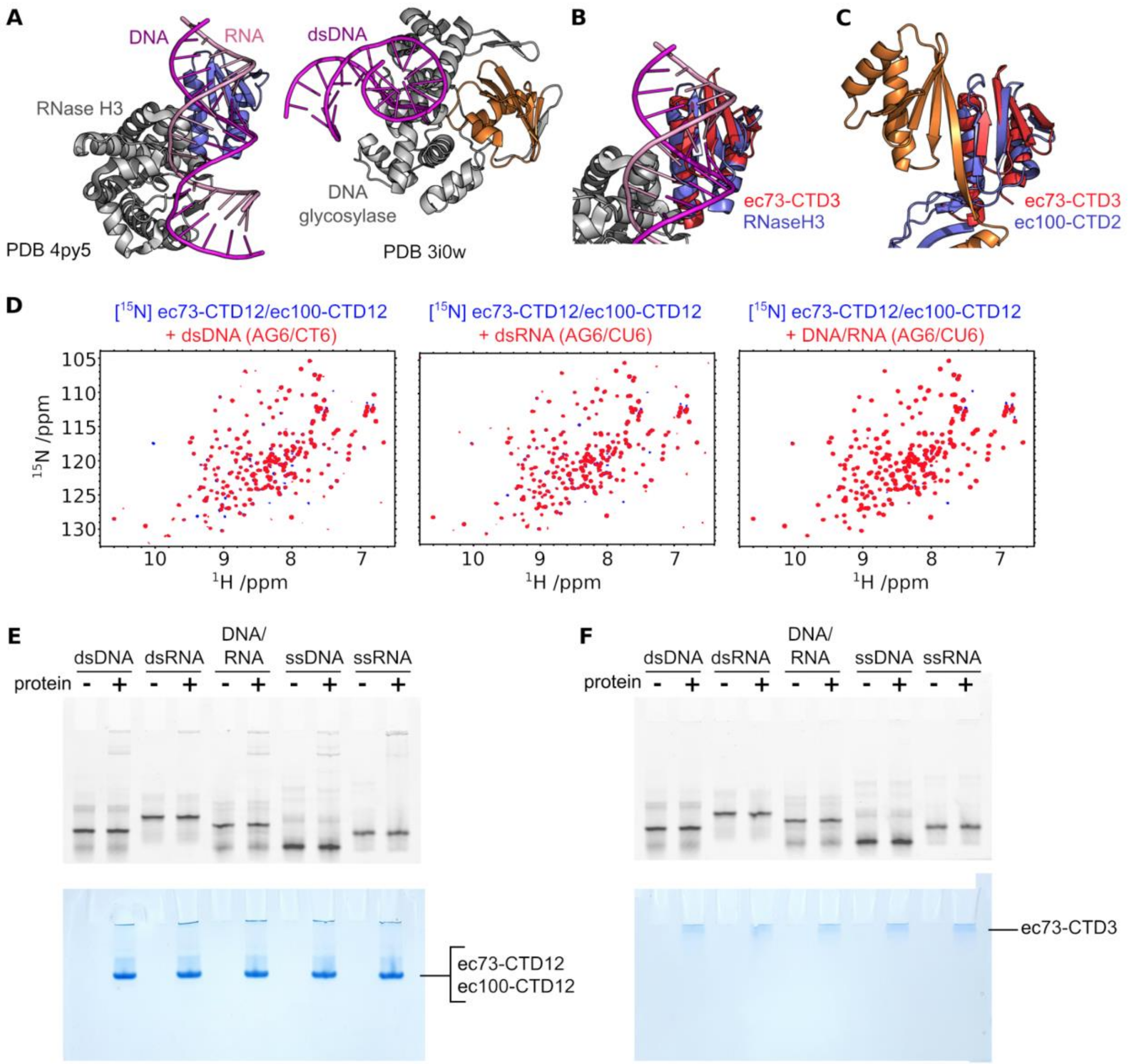


**Supplementary Figure S4**. The TBP domains from ecCPSF73 and ecCPSF100 do not bind nucleic acids. (A) Four-stranded TBP domain (violet) located in archaeal RNase H3 bound to a DNA-RNA duplex (PDB ID 4py5)((50), and the extended TBP domain (orange) from *Clostridium* DNA glycosidase bound to dsDNA (PDB ID 3i0w)(62). (**B**) Superposition of ec73-CTD3 (red) with RNaseH3 from panel (A). Calculated RMSD of 2.5 Å. (**C**) Superposition of ec73-CTD3 with ec100-CTD2. (**D**) $^{15}$N-HSQC spectra for unbound 100 μM [$^{15}$N]ec73-CTD12/ec100-CTD12 (blue spectra), overlaid with the spectra of the same samples but in the presence of 150 μM dsDNA (red spectrum, left), 150 μM dsRNA (red spectrum, middle) or 150 μM DNA/RNA duplex (red spectrum, right). At these concentrations, even relatively weak interactions would be apparent. (**E**) Native gel EMSA to assay nucleic acid binding, using 500 nM Cy3-labelled dsDNA, dsRNA, DNA/RNA duplex, ssDNA, or ssRNA, in the absence or presence of 50 μM ec73-CTD12/ec100-CTD12. The top image uses fluorescence to image the nucleic acids, and the bottom image is the same gel following Coomassie staining to image the proteins. (**F**) The same set-up as for (E) but using ec73-CTD3.

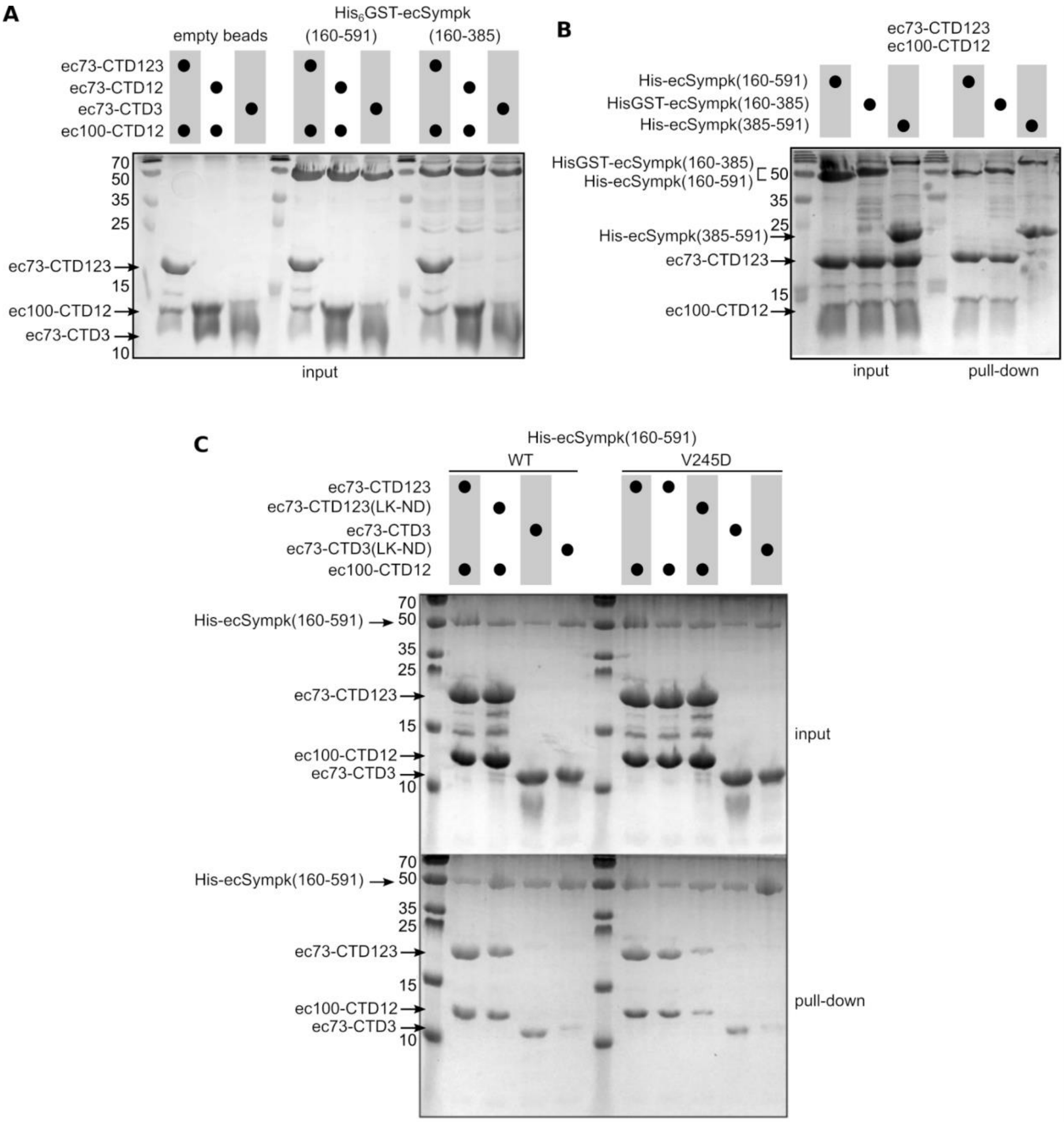


**Supplementary Figure S5**. Additional SDS-PAGE gels related to the pulldown experiments in Figures 5 and 6. (**A**) SDS-PAGE gel of the sample inputs from the experiment shown in **Figure 5D**. (**B**) SDS-PAGE results from a pulldown experiment showing that the ec73-CTD123/ec100-CTD12 complex binds to ecSympk residues 160-385, and not to residues 385-591. (**C**) SDS-PAGE results from additional samples related to the experiments shown in **Figure 6C**.

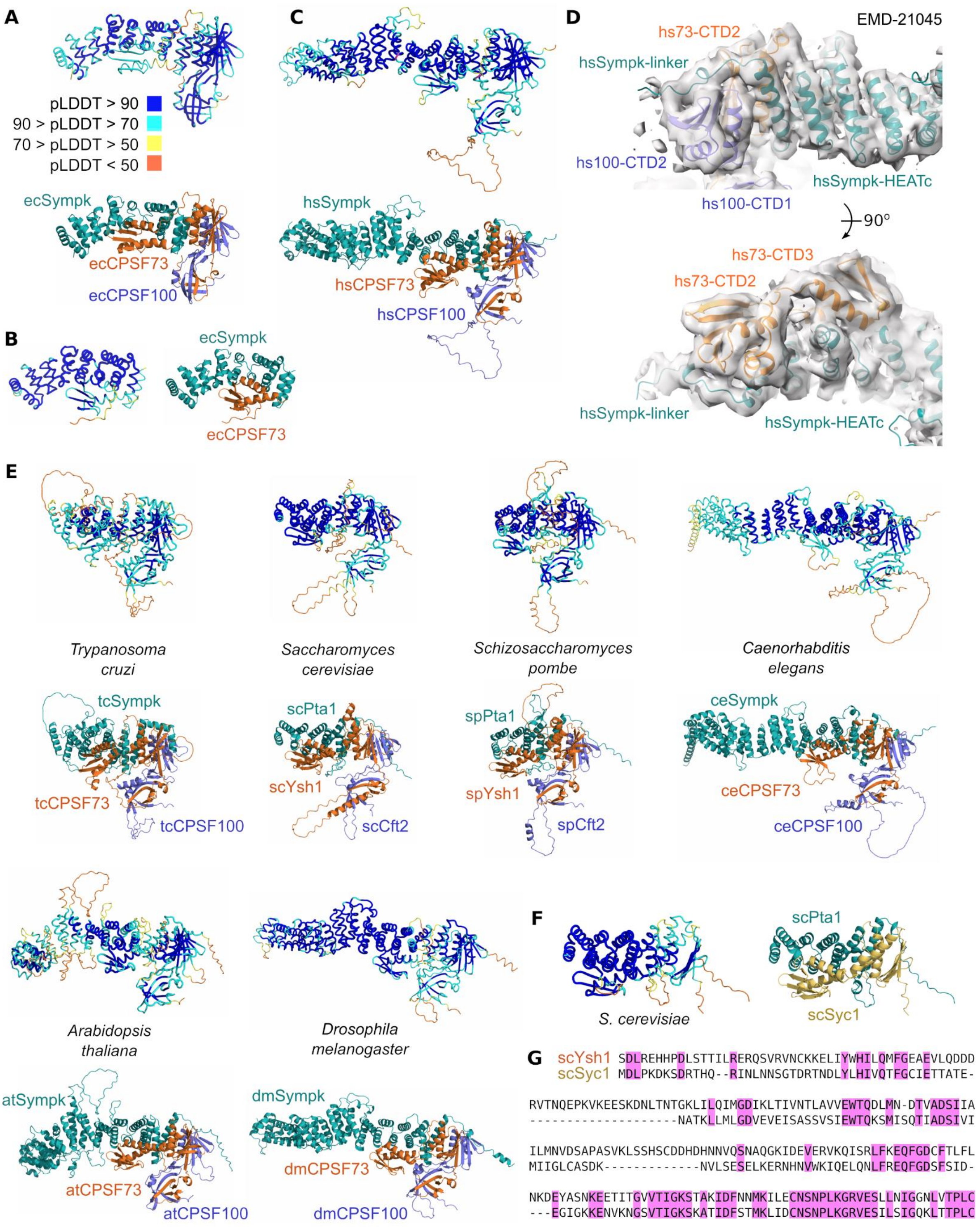


**Supplementary Figure S6**. AlphaFold2 models of C-terminal CPSF73-CPSF100-Sympk trimer complexes. (**A**) Top pTMscore-ranked AlphaFold2 model of the *E. cuniculi* CPSF73-CPSF100-Sympk C-terminal complex, shown in tubes coloured by pLDDT (top), and as cartoon coloured by chain (bottom). (**B**) Similar representation of the AlphaFold2 model using just sequences from ec73-CTD3 and ecSympk(160-385). (**C**) Top pTMscore-ranked AlphaFold2 model of the human CPSF73-CPSF100-Sympk C-terminal complex. (**D**) AlphaFold2 model of the human trimer (C) docked into the cryo-EM map of the histone pre-mRNA 3'-end processing complex (EMD-21045)(Sun et al., 2020). (**E**) Top pTMscore-ranked AlphaFold2 models from other species. (**F**) Top pTMscore-ranked AlphaFold2 model of the complex between *S. cerevisiae* Syc1 and the C-terminus of Pta1. (G) Alignment of the CTD2-CTD3 region of *S. cerevisiae* Ysh1 with full-length Syc1. Identical residues are highlighted in magenta, used for **Figure 7C**.
